# Zebrafish lack a strong meiotic checkpoint response to defects in chromosome synapsis

**DOI:** 10.1101/2025.03.18.644038

**Authors:** Iván Olaya, Ilara N. Yilmaz, Naima Nour-Kasally, Bruce W. Draper, Sean M. Burgess

## Abstract

The synaptonemal complex (SC) is a meiosis-specific structure that aligns homologous chromosomes and facilitates the repair of meiotic DNA double-strand breaks (DSBs). Defects in SC assembly or unrepaired DSBs trigger a prophase I checkpoint to prevent the formation of aneuploid gametes. The strength of these checkpoints varies among species and between sexes. Whether zebrafish (*Danio rerio*) have prophase I surveillance mechanisms that monitor chromosome synapsis and/or meiotic DSB repair has not been explored. To investigate how defects in SC formation affect gametogenesis in zebrafish, mutations in genes encoding two structural components of the SC, *syce2* and *sycp1* were examined. While *syce2* and *sycp1* fish exhibit defects in both synapsis and DSB repair, the two mutants show different reproductive outcomes. *syce2* mutant females and males produce a significant percentage of normal progeny. In contrast, *sycp1* mutant females produce fewer normal offspring, while *sycp1* mutant males are infertile, with spermatocytes arrested at metaphase I. Notably, offspring from *syce2* and *sycp1* mutant mothers show extensive somatic mosaic aneuploidy, indicating that defects in the meiotic machinery can lead to genome instability during embryogenesis. Our findings suggest that a checkpoint monitoring the progression of synapsis may be weak or absent in the zebrafish, resembling the situation in yeast, plants, and *Drosophila*, rather than in mice and the nematode *Caenorhabditis. elegans* where robust mechanisms exists to silence unpaired chromosomes leading to meiocyte apoptosis.

**Author Summary:** Meiosis is a specialized cell division that reduces chromosome numbers for the production of haploid gametes and is necessary for sexual reproduction. During meiosis prophase I, precise coordination is required for key events such as the formation and repair of DNA double-strand breaks and the synapsis of homologous chromosomes, which is mediated by the synaptonemal complex. Specialized checkpoint pathways monitor the proper execution of these events. Errors in either process can induce a checkpoint response leading to cell death or the production of aneuploid gametes. Here, we demonstrate that the checkpoint monitoring synapsis is weak—or potentially absent—in zebrafish, suggesting that a robust synapsis checkpoint is not conserved across all vertebrates. We discuss this finding in the context of the teleost-specific whole genome duplication and the widespread heterogametic switching (e.g., XY to ZW systems) within this diverse clade, which includes over 30,000 species.

## Introduction

There has been a world-wide trend of decreasing sperm count among men, yet the cause is unknown [1]. Mutations in genes encoding protein components of the meiotic synaptonemal complex (SC) are associated with nonobstructive azoospermia [2–5] and pregnancy loss [6] and primary ovarian insufficiency syndrome in women [7,8]. It is possible that effects of less deleterious mutations might be enhanced by environmental factors [1]. Studies in several model organisms have demonstrated the critical role the SC plays in the completion of meiosis that have provided insights into human infertility [9].

Meiosis is a specialized cellular program that is required to produce haploid gametes through one round of DNA replication and two consecutive rounds of chromosome segregation. Meiosis I segregates homologous chromosomes (homologs) while meiosis II separates sister chromatids. Crossing over between non-sister chromatids of homologs by homologous recombination is central to proper meiosis I segregation. Recombination is initiated by forming and repairing programmed DNA double-strand breaks (DSBs) that are catalyzed by the enzyme Spo11 [10,11]. Each homolog pair experiences at least one crossover that supports proper segregation [12–17]. This so-called “crossover assurance” depends on ZMM proteins that function within the context of the synaptonemal complex (SC) [18,19].

The SC is an evolutionarily conserved tripartite structure observed in meiotic prophase across many species [20] and is composed of two chromosome axes, one for each homolog, a central region containing proteins that connect homologous axes, and a central element within the central region that stabilizes the overall structure (Fig 1A) [21,22]. In vertebrates, the central region contains head-to-head interacting dimers of the transverse filament protein SYCP1 flanked by two bands of central element filaments containing SYCE1, SYCE2, SYCE3, TEX12 and SIX6OS1 [21]. In mouse, SYCE2 and TEX12 form a complex that stabilizes interactions between the αN domains of SYCP1 [23–27]. Once the SC is established at sites of future crossovers, the central element filaments promote expansion of the SC to align chromosomes end-to-end. Mouse SYCE2 is required to fully assemble the SC by stabilizing SYCP1 interactions, and its absence leads to foci or short stretches of SYCP1 between paired chromosome axes [23]. Functional orthologs of the central element proteins also exist in the SC in budding yeast [28] and *Drosophila* [29], highlighting the conserved nature of SC assembly.

**Figure 1.**
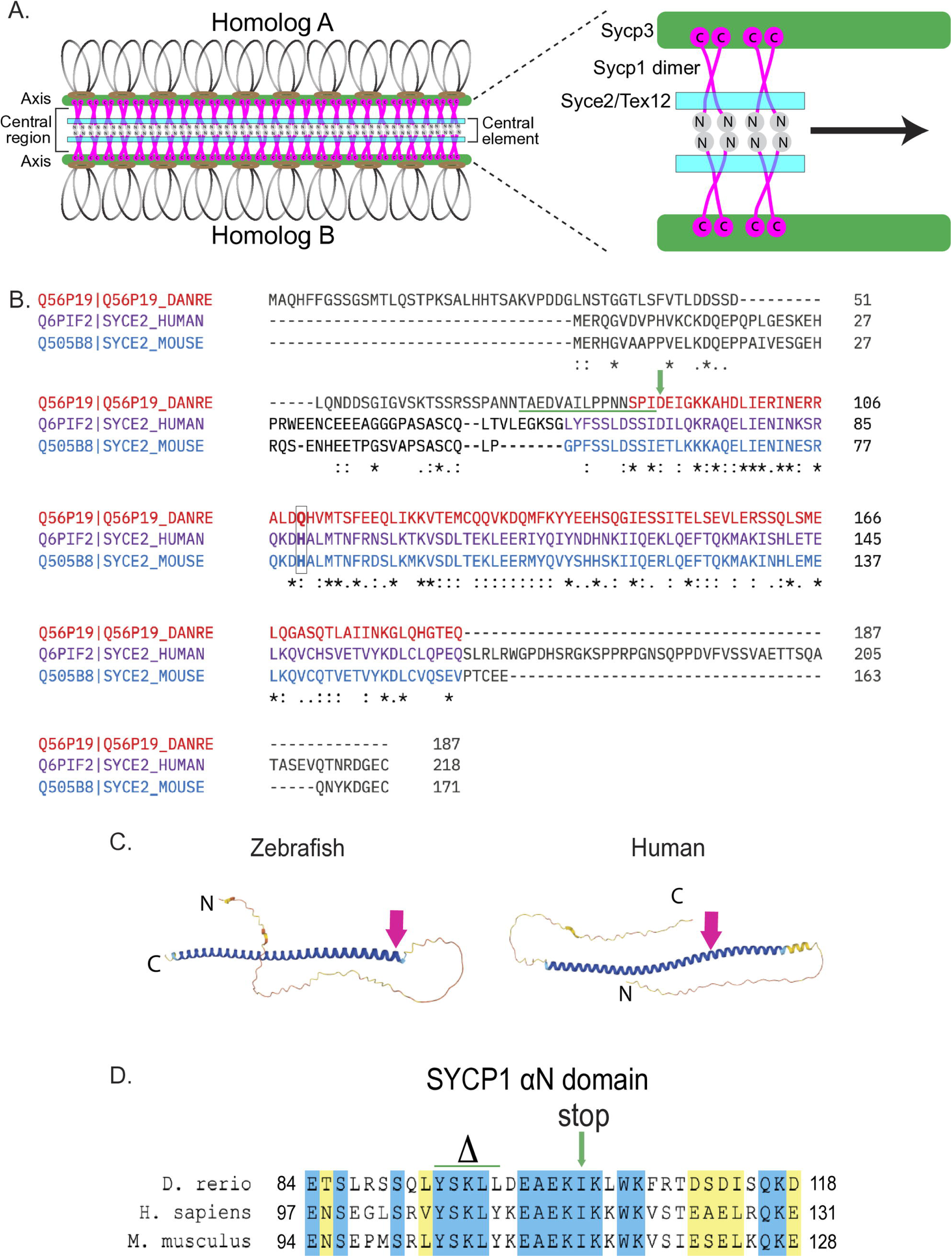
Generating the *syce2* and *sycp1* mutants. (A) Schematic of the synaptonemal complex showing key structures of meiotic chromosomes: DNA loops (gray/black), chromosome axis (Sycp3; green), central region (Sycp1; magenta) and the central element (Syce2/Tex12; cyan) that includes the Sycp1 N-termini (gray). Other known central element components not shown: Syce1, Syce3 and Six6os1. (B) Alignment of the zebrafish Syce2 amino acid sequence with human and mouse orthologs using the Clustal Omega multiple sequence alignment tool [129]. Colored text defines regions that form a long α helix in a 2:2 complex [26]. Conserved identical residues (*), strongly similar (:) and weakly similar (.). The amino acids (aa) affected by the frameshift mutation are underlined in green. The position of the premature stop codon in *syce2^−/−^* is shown by a green arrow. The relative position of the human mutation (H89Y) associated with pregnancy loss [6] is highlighted in the boxed region. (C) AlphaFold-predicted structures for Syce2 protein from zebrafish (AF-Q56P19-F1) and human (AF-Q6PIF2-F1) [131]. The position of the 89th codon for zebrafish Syce2 that generates the premature stop codon is shown by a pink arrow. The position of the human variant in SYCE2 is depicted by the pink arrow. (D) Alignment of aa comprising the αN domain of Human SYCP1 that promotes self-assembly [25]. Blue regions are identical aa and yellow regions are similar aa. The CRISPR generated *sycp1* mutation is a complex mutation with a deletion that removes the amino acids marked by delta and an insertion that truncates the protein at the amino acid marked by the arrow. See Fig S1 for details.

A meiotic checkpoint network exists to coordinate the ordered execution of dependent events of meiotic prophase I [30]. Asynapsis or unrepaired DSBs can lead to delayed or arrested prophase I, and in some cases result in apoptosis. In mice and the nematode *C. elegans*, the failure to fully synapse homologs activates a checkpoint that is separable from one surveilling DSBs, and in both cases cells are eliminated at meiotic prophase I [31–38]. By contrast, meiotic checkpoints that monitor synapsis progression are relatively weak, or absent, in budding yeast and female *Drosophila* and in *Arabidopsis*. In yeast, disrupting the transverse filament or central element delays the first meiotic division, presumably due to the slow repair of DSBs and not through a synapsis checkpoint *per se* [39–43]. In *Drosophila* females, partial disruption of the SC leads to a checkpoint-mediated prophase I delay, yet total loss of SC does not appear to cause delay or arrest [44,45]. In *Arabidopsis*, elimination of the SC has marginal effects on seed number and pollen viability [46]. Zebrafish *spo11* mutants fail to initiate homologous recombination and synapsis, yet females produce normal numbers of fertile eggs [47]. This observation raises the question if a meiotic checkpoint monitoring synapsis operates in zebrafish. If not, this would demonstrate that a synapsis checkpoint is not a general feature of meiosis in vertebrates. Since Spo11-dependent DSBs and synaptonemal complex are absent in this mutant, it is possible that no “checkpoint-inducing” signal is being produced, similar to what has been proposed for the *spo11* mutant in budding yeast [48,49]. In this study, we analyzed *syce2* and *sycp1* mutants that exhibit aberrant SC formation and/or inefficient repair of DSBs.

The chromosome events of meiotic prophase I follow a coherent timeline in zebrafish [47,50]. During leptotene, the deposition of short axial structures at each telomere is concurrent with the formation of DSBs. At early zygotene, co-alignment of short homolog axes is followed by the initiation of the SC, which extends closely behind the growing axes creating a moving fork until homologs are synapsed from end-to-end at pachytene. Co-alignment and synapsis appear to initiate almost exclusively at the ends of chromosomes while interstitial regions are joined by “zippering up”. This is reminiscent of the temporal order of events in *C. elegans*, where pairing and synapsis are initiated solely at defined pairing centers located at the ends of chromosomes and the SC extends in a linear fashion [37,51,52].The important difference is that in *C. elegans*, pairing at chromosome ends is mediated by protein/protein interactions to bridge homologs, while in zebrafish it is mediated by the processing of Spo11-induced DSBs.

A mutation in the zebrafish *sycp1* gene was recovered in a forward genetic screen to identify mutants defective for spermatogenesis [53]. Further analysis of mutant *sycp1^isa^*^/isa^ males showed that homologs pair at the ends but do not synapse [54]. In addition, spermatogenesis was arrested, though the meiotic stage of the arrest was not determined. Since females were not analyzed in this study, it remained unknown how loss of *Sycp1* affects oogenesis. Here, we examined fish containing loss-of-function *syce2^−/−^* and *sycp1^−/−^* mutations and found that synapsis is completely abolished in *sycp1^−/−^* mutants and reduced to short synapsed-like structures at the ends of chromosomes in *syce2^−/−^* mutants. Both mutants exhibit increased numbers of DSBs in zygotene nuclei and inefficient repair implicating the SC in this process. Remarkably, *syce2^−/−^* mutant males and females are fertile and produce a high percentage of healthy offspring. By contrast, *sycp1^−/−^* males are infertile with spermatocytes arresting at metaphase I. In addition, *sycp1^−/−^* females produce predominantly malformed progeny, though some live to adulthood. These data show that short stretches of SC are sufficient to produce normal progeny and that asynapsis or unrepaired DSBs in zebrafish do not elicit a strong, if any, prophase I checkpoint response. Our study sheds light on the stringency of meiotic checkpoints and highlights the differences in reproductive outcomes between males and females in a teleost species. We also show that embryos of *syce2^−/−^* and *sycp1^−/−^* mothers exhibit a striking degree of somatic mosaic aneuploidy suggesting that defects in meiotic genes can lead to genome instability during embryogenesis.

## Results

### Targeted mutation of *syce2* and *sycp1*

The zebrafish *syce2* gene consists of 5 exons that encode a 187 amino acid (aa) protein. Alignment of the zebrafish Syce2 amino acid sequence with human and mouse orthologs highlights conservation of the core region of SYCE2 in vertebrates (Fig 1B). The core is a long α helical structure shown to interact with TEX12 [24]. A human *SYCE2* genetic variant associated with pregnancy loss is located within the core region [6] (Fig 1B). To disrupt the function of zebrafish *syce2*, we used CRISPR-Cas9 to introduce an indel mutation resulting in a predicted 88 aa truncated protein (*syce2^uc98^*; Fig 1B; Supplemental Fig S1A). An Alphafold predicted structure shows that the *syce2^uc98^* mutation should remove all but three amino acids of the core and abolish the Tex12 interaction domain (Fig 1C). Antibodies to mammalian SYCE2 did not cross-react with the zebrafish protein. In this study we refer to the *syce2^uc98/uc98^* mutant as *syce2*^−/−^.

The *sycp1* gene contains 32 exons that encode a 1000 aa protein. Using CRISPR-Cas9 we introduced a deletion/insertion mutation into exon 5 (*sycp1^uc97^*; Supplemental Fig S1B). This mutation creates a net five aa deletion within the αN domain immediately followed by a premature stop codon resulting in a predicted 97 aa protein (Fig 1D). Sequencing of RT-PCR products generated from *syce2^uc98^* and *sycp1^uc97^* testes RNA confirmed that the mutations do not cause exon skipping that could potentially lead to a truncated but functional protein (Supplemental Fig S1C) [55,56]. For this study we refer to the *sycp1^uc97/uc97^* mutant as *sycp1^−/−^.* Immunostaining Sycp1 reveals its localization is absent in *sycp1^−/−^* mutants (described below).

### Loss of Syce2 leads to asynapsis of interstitial regions of homologous chromosomes

Since mammalian SYCE2 functions to extend the SC from sites of initiation [23], we predicted that in *syce2^−/−^* zebrafish, synapsis would occur at the ends of chromosomes but would fail to “zipper-up” end-to-end. To test this, we analyzed images of immunostained surface-spread chromosomes from wild-type and *syce2^−/−^* zebrafish spermatocytes using structured-illumination microscopy. In wild-type spermatocytes, the homolog axes assemble initially at the chromosome ends in leptotene, co-align in early zygotene, then extend concurrently with the deposition of Sycp1 to form full-length SC in pachytene (Fig 2A). In *syce2^−/−^* spermatocytes, the majority of the 50 chromosome ends form short parallel axial tracks in early zygotene, suggesting that homologs have found their partners (Fig 2B). However, the synapsed-like configuration at the ends does not lengthen as meiosis prophase progresses. Instead, the axes continue to elongate until they include the entire length of each homolog, but the interstitial regions remain separated. Occasional patches of synapsed-like axes occur in the interstitial regions in the *syce2^−/−^* mutant but are relatively rare. We measured the mean lengths of synapsed and synapsed-like segments per nucleus which are 7.71 ± 1.54 µm in wild-type pachytene nuclei compared to 1.16 ± 0.26 µm in pachytene-like nuclei of the mutant (Fig 2C). While the chromosomes in the *syce2^−/−^* spreads were too entangled to allow each chromosome to be traced from end to end, we estimate that ~ 30% of the total chromosome length in *syce2^−/−^* mutants is in a synapsed-like configuration.

**Figure 2.**
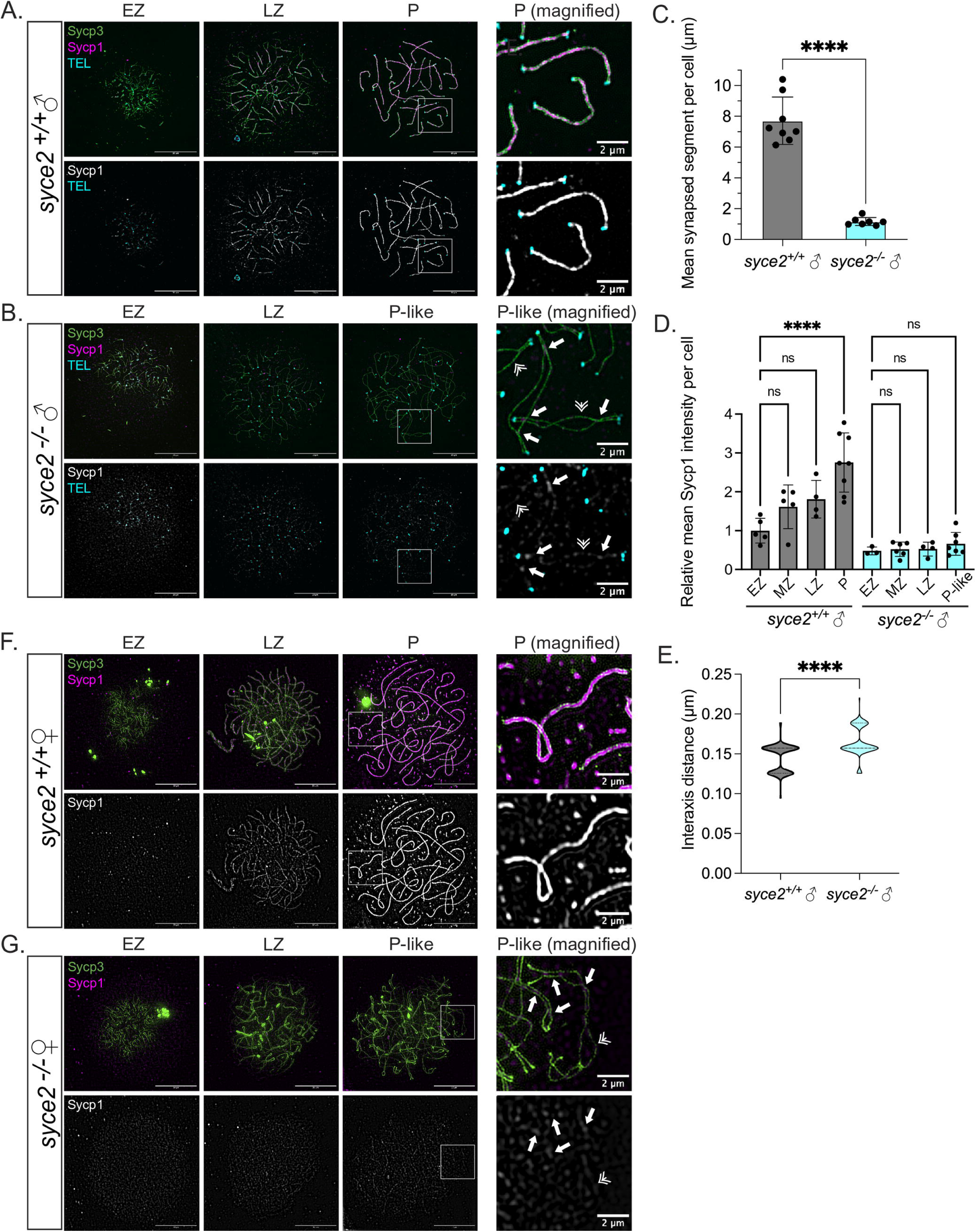
Loss of Syce2 function results in asynapsed interstitial regions. (A–B) Surface-spread chromosomes from *syce2*^+/+^ (A) and *sycpe2^−/−^* (B) spermatocytes imaged using structured illumination microscopy. Proteins are immunostained for Sycp3 (green) and Sycp1 (magenta and gray). Telomeres (cyan) are detected by *in-situ* hybridization using a fluorescent PNA probe. Examples of spread chromosomes during meiotic prophase are shown: Early Zygotene (EZ), Late Zygotene (LZ), Pachytene (P) (*syce2*^+/+^), and Pachytene-like (P-like) (*sycpe2^−/−^*). Scale bar = 10 µm. Boxed region represents magnified examples of synapsed pachytene (*syce2*^+/+^) and pachytene-like (*syce2^−/−^*) chromosomes. Scale bar for magnified examples = 2 µm. Arrows represent synapsed-like regions with Sycp1 localization. Double arrowheads indicate Sycp1 localization at unaligned axial regions. (C) Mean synapsed segment per cell (µm) in pachytene and pachytene-like spermatocytes. The mean synapsed segment is significantly lower in *syce2*^−/−^ spermatocytes (n = 7) compared to *syce2*^+/+^ (n = 8). Significance was determined using an unpaired t-test. **** = p < 0.0001. (D) Relative mean Sycp1 intensity per cell from early zygotene to pachytene or pachytene-like spermatocytes normalized to the mean *syce2*^+/+^ EZ values. Mean-fluorescence intensity between the axes (n=25) – mean background (n=6) was determined for each cell using the segmented-line tool in ImageJ. n = 5, *syce2*^+/+^ – EZ; n = 5; *syce2*^+/+^ – MZ; n = 4, *syce2*^+/+^ – LZ; n = 8, *syce2*^+/+^ – P; n = 3, *syce2*^−/−^ – EZ; n = 6, *syce2*^−/−^ – MZ; n = 4, *syce2*^−/−^ – LZ; n = 7, *syce2*^−/−^ – P-like. Significance was determined using ordinary one-way ANOVA testing with Šidák’s multiple comparisons test. ns = not significant; **** = p < 0.0001. (E) Distance between axes for co-aligned axial pairs in pachytene (n = 126) and pachytene-like (n = 93) spermatocytes from 8 cells in *syce2*^+/+^ and 7 cells in *syce2^−/−^* shown as a violin plot. Note that one pixel is equal to 0.03 µm resulting in a graph with a bimodal distribution. Significance was determined using an unpaired t-test. **** = p < 0.0001. (F–G) Surface-spread chromosomes from *syce2*^+/+^ (G) and *sycpe2^−/−^* (H) oocytes imaged using structured illumination microscopy immunostained for Sycp3 and Sycp1. Examples of spread chromosomes during meiotic prophase are described in (A–B). Scale bar = 10 µm. Boxed region represents magnified examples of synapsed pachytene and pachytene-like chromosomes. Scale bar for magnified examples = 2 µm. Arrows represent synapsed-like regions with Sycp1 localization. Double arrowhead indicates Sycp1 localization at unaligned axial regions.

To better understand how synapsed-like regions compare to *bona fide* SC, we measured the relative levels of Sycp1 protein in synapsed (wild-type) and synapsed-like (*syce2^−/−^*) chromosomes in nuclei that represented all stages of prophase. In wild-type spermatocytes, the relative mean Sycp1 fluorescence intensity increased steadily throughout prophase I from 1.00 ± 0.32 in early zygotene to 2.75 ± 0.76 in pachytene (Fig 2D). In *syce2^−/−^* spermatocytes, Sycp1 also loads between nascent axes, but at levels barely above background, and in foci or short patches instead of continuous stretches (Fig 2B; P-like magnified). Sycp1 is also found at low levels along the lengths of asynapsed axes. In addition, the relative mean intensity of Sycp1 between axes in *syce2^−/−^* spermatocytes remains low from early zygotene (0.48 ± 0.10) through the pachytene-like stage (0.66 ± 0.29) (Fig 2D). The diminished Sycp1 levels observed in the *syce2^−/−^* mutant presumably reflect the destabilization of the central region.

We next tested whether the reduced level of Sycp1 on *syce2^−/−^* chromosomes affects the distance between axes in the synapsed-like regions. For this, a line perpendicular to the parallel axes was drawn in ImageJ and the distance between the maximum intensity signal (peaks) for each axis was determined. We found that the mean distance between peaks was slightly larger in the *syce2^−/−^* mutant (0.17 ± 0.02 µm) compared to wild-type (0.14 ± 0.02 µm) (Fig 2E). These results show that the synapsed-like regions in *syce2^−/−^* mutants resemble wider than normal SC with reduced amounts of Sycp1.

We next examined chromosome spreads of *syce2^+/+^* and *syce2^−/−^* oocytes (Fig 2F and G). As seen for spermatocytes, Sycp1 also appears as foci at sub-telomeric regions in *syce2^−/−^* zygotene oocytes and as short stretches in pachytene-like oocytes (Fig 2G). While we did not observe full-length chromosome alignment in the *syce2^−/−^* females, the co-aligned (< 0.5 µm) and synapsed-like regions are found both at the ends of chromosomes and in interstitial regions. Since the distribution of Mlh1 foci is more evenly distributed in zebrafish oocytes compared to spermatocytes [57,58], it appears that partial synapsis may mirror the different recombination landscape of the two sexes. Notably, there was considerable entanglement of axes in both oocytes and spermatocytes.

### Co-aligned axes but not synapsis in *sycp1^−/−^* oocytes

A previous study of *sycp1^isa^*^/isa^ failed to recover females in the homozygous population [54], therefore how loss of Sycp1 affects oogenesis was not determined. Because we found females among the *sycp1^−/−^* population, we could examine chromosome spreads of *sycp1^−/−^* females. We found evidence of chromosome co-alignment without synapsis at sub-telomeric regions in early zygotene as in wild-type (Fig 3A and Fig 2F), but full-length alignment is not observed in later stage oocytes. These results show that synapsis is required for end-to-end alignment of homologs during oogenesis.

**Figure 3.**
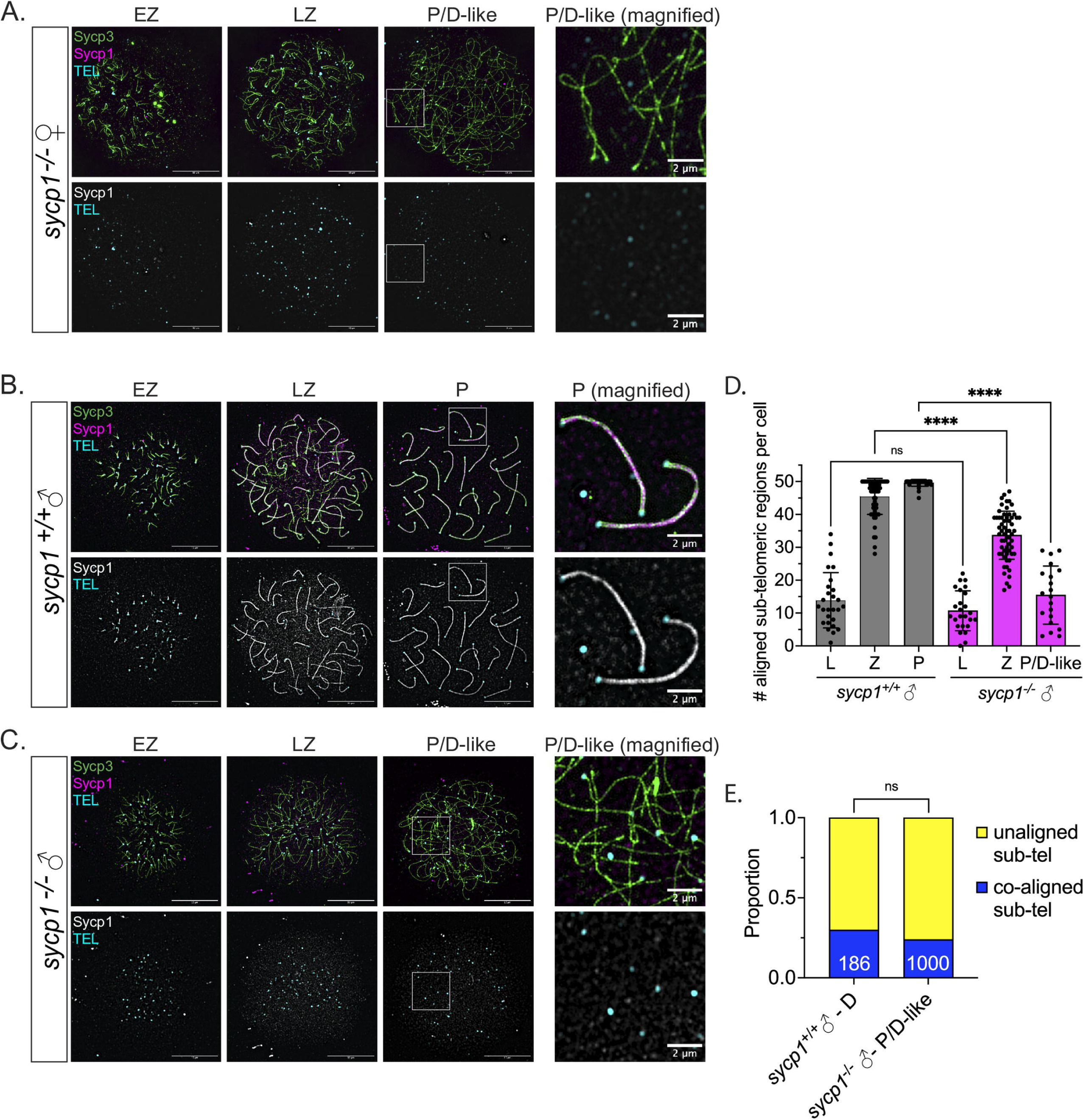
Formation and loss of co-alignment in *sycp1^−/−^* spermatocytes. (A) Surface-spread chromosomes from *sycp1^−/−^* oocytes imaged using structured illumination microscopy stained for Sycp3 (green), Sycp1 (magenta and gray) and telomeres (cyan). Examples of spread chromosomes during meiotic prophase are shown: EZ, LZ, P and pachytene/diplotene-like (P/D-like) (*sycp1^−/−^*). Scale bar = 10 µm. Boxed region represents magnified examples of P/D-like chromosomes. Scale bar for magnified examples = 2 µm. (B-C) Surface-spread chromosomes from *sycp1*^+/+^ (A) and *sycp1^−/−^* (B) spermatocytes imaged using structured illumination microscopy stained for Sycp3 (green), Sycp1 (magenta and gray) and telomeres (cyan). Examples of spread chromosomes during meiotic prophase are described in (A). Scale bar = 10 µm. Boxed region represents magnified examples of P and P/D-like chromosomes (*syce2*^+/+^). Scale bar for magnified examples = 2 µm. (D) Number of co-aligned pairs of sub-telomeric regions per cell across prophase in *sycp1*^+/+^ and *sycp1*^−/−^ spermatocytes from (A–B). Images for leptotene (L) are not shown but are characterized by predominantly short un-aligned axes. n = 27, *sycp1*^+/+^– L; n = 25, *sycp1*^−/−^– L; n, = 61 *sycp1*^+/+^– Z; n = 60, *sycp1*^−/−^– Z; n = 36, *sycp1*^+/+^– P; n = 20, *sycp1*^−/−^– P/D-like. Significance was determined using ordinary one-way ANOVA testing with Šidák’s multiple comparisons test. ns = not significant; **** = p < 0.0001. (E) Proportion of co-aligned and unaligned pairs of sub-telomeric regions in *sycp1*^+/+^ diplotene (D) (n = 186) and *sycp1*^−/−^ P/D-like (n = 1000) spermatocytes. 11 cells were used for *sycp1*^+/+^ and 20 cells for *sycp1*^−/−^. Significance was determined using Fisher’s exact test. ns = not significant.

### Formation and loss of co-aligned axes in *sycp1^−/−^* spermatocytes resembles wild-type chromosome behavior

A previous study of the *sycp1^isa^* allele reported that sub-telomeric regions of *sycp1^isa^*^/isa^ spermatocytes were aligned in leptotene and early zygotene, but that these tight associations were lost at late prophase stages [54]. We also found co-aligned ends during early zygotene (EZ) and late zygotene (LZ) stage of spermatogenesis that were similar to wild-type chromosome spreads (Fig 3B and Fig 3C). We found the number of co-aligned pairs of chromosome ends per cell in *sycp1*^−/−^ leptotene spermatocytes (10.7 ± 6.1) out of 50 is comparable to wild-type (13.9 ± 8.4) (Fig 3D) and these values increase in zygotene (33.7 ± 7.3), yet they do not reach the same level as synapsed ends in wild-type (45.5 ± 5.5). Instead, the number of co-aligned regions in pachytene/diplotene-like *sycp1^−/−^* mutant spermatocytes is reduced from zygotene levels by ~54% (15.5 ± 8.8) consistent with the *sycp1^isa^*^/isa^ phenotype which was attributed to possibly weak contacts at sub-telomeric regions or DSBs resolving into non-crossovers [54]. Similar to *syce2^−/−^* mutants, there is considerable entanglement of axes in both oocytes and spermatocytes.

We next considered that the loss of co-aligned ends in *sycp1^−/−^* pachytene/diplotene might reflect a natural feature of diplotene chromosomes. Diplotene is the stage following pachytene when the SC is disassembled, and homologs are held together by chiasmata (sites of crossing over). Thus, the proximity of telomeres in diplotene bivalents depends on the distance between a chiasma and the adjacent chromosome ends. Since zebrafish spermatocytes experience ~1.1 crossovers per chromosome on average [57,59], it would be expected that most telomeres of diplotene chromosomes will not be in close proximity. We set out to measure the proportion of nearby ends separated by a distance < 0.5 µm in *sycp1^−/−^* mutant spreads with full-length axes and in wild-type diplotene chromosomes. Since asynapsis of wild-type diplotene chromosomes is not uniform across all chromosomes in a given nucleus, we limited our analysis to bivalents with full-length axes and large scale asynapsis (Fig S2). The proportion of unsynapsed aligned ends in wild-type diplotene spermatocytes (29.6%) is indistinguishable from *sycp1^−/−^* pachytene/diplotene-like spermatocytes (24.2%) (Fig 3E). Therefore, loss of sub-telomeric contacts is a general property of normal diplotene chromosomes and *sycp1^−/−^* chromosomes with full-length axes may be in a diplotene-like configuration.

#### Different reproductive outcomes in *syce2^−/−^* and *sycp1^−/−^* mutants despite severe synapsis defects

In mice, mutations in *Syce2* and *Sycp1* lead to infertility due to a prophase checkpoint that removes defective spermatocytes and oocytes [21]. We therefore tested whether fertility was affected in *syce2^−/−^* and *sycp1^−/−^* mutants. *syce2^−/−^* males induce wild-type females to spawn similar numbers of eggs (272.4 ± 164.9) as wild-type males (234.5 ± 103.6) (Fig 4A), indicating normal mating behavior. When crossed to wild-type males, *syce2^−/−^*females also yield similar numbers of spawned eggs (183.6 ± 101.0) as wild-type females (207.6 ± 146.4) (Fig 4A). Average fertilization rates of *syce2^−/−^* males (74.6 % ± 26.4%) and females (95.2 % ± 4.8%), respectively, are similar to wild-type controls (90.4 % ± 9.2% and 90.1 % ± 25.7%, respectively) (Fig 4A). These data show that both male and female *syce2^−/−^* mutants are fertile, despite severe defects in synapsis, suggesting that a strong prophase checkpoint monitoring synapsis is not activated in this species.

**Figure 4.**
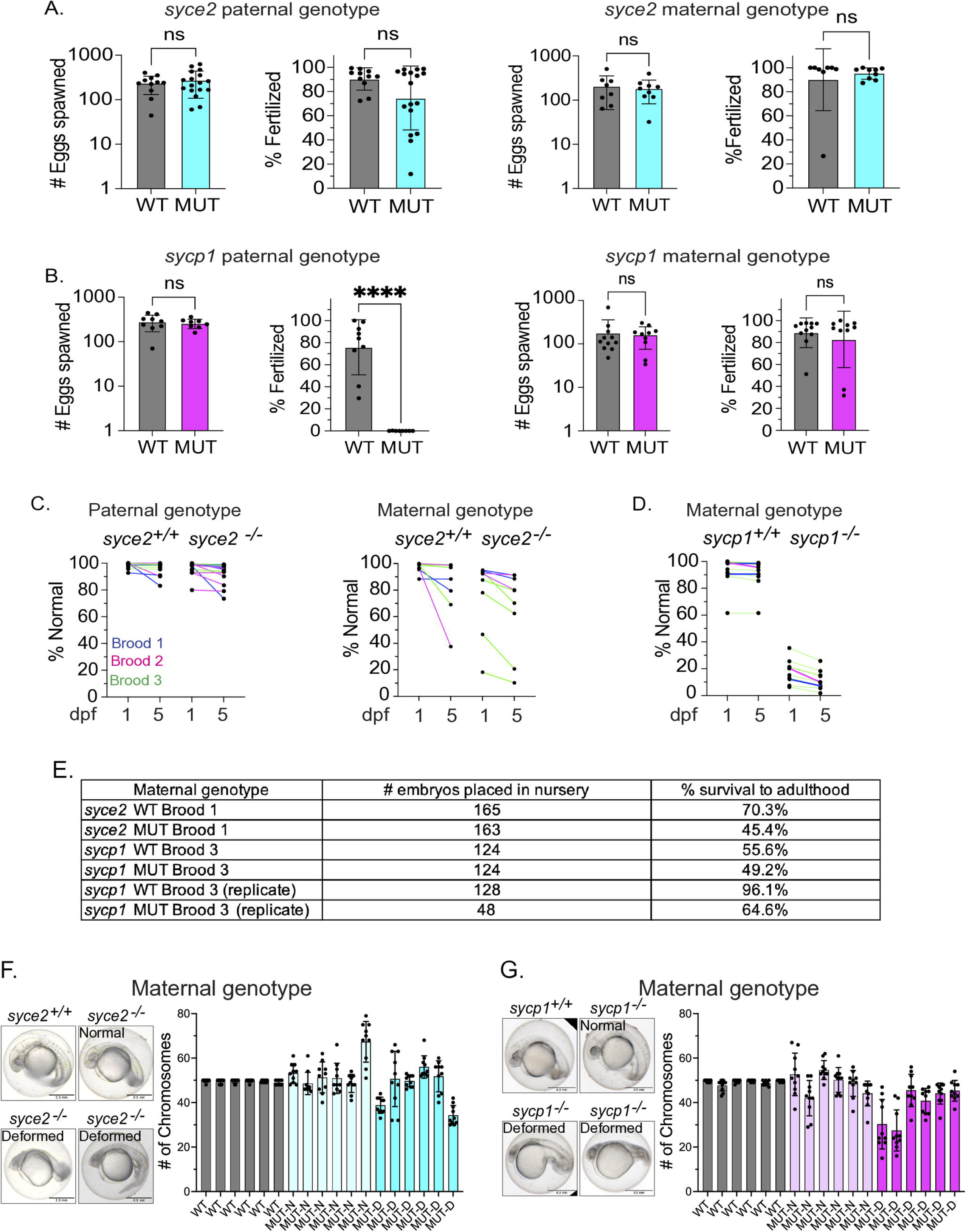
*syce2^−/−^* and *sycp1^−/−^* mutants have different reproductive outcomes. (A) Number of eggs spawned and fertilization rate per cross from wild-type and *syce2*^−/−^ mutants. Each data point represents a cross yielding > 20 eggs from individual animals from three different broods born on different days. Percent fertilized excludes decomposed eggs. *syce2*^+/+^ males, n = 11; *syce2*^−/−^ males n = 17; *syce2*^+/+^ females n = 8; and *syce2*^−/−^ females, n = 9. Significance was determined using an unpaired t-test. Ns = not significant. (B) Number of eggs spawned and fertilization rate per cross for wild-type and *sycp1*^−/−^ mutants. Each data point represents a cross yielding > 20 eggs from individual animals from three different broods born on different days. Percent fertilized excludes decomposed eggs *sycp1*^+/+^ males, n = 9; *sycp1*^−/−^ males n = 8; *sycp1*^+/+^ females n = 11; and *sycp1*^−/−^ females, n = 10. Significance was determined using an unpaired t-test. ns = not significant.; **** = p < 0.0001. (C) Percent Normal-looking offspring at 1- and 5-days post fertilization (dpf) from wild-type and *syce2* mutants from data in (A). Colored lines represent animals from 3 broods. (D) Percent Normal-looking offspring at 1 and 5 dpf from wild-type and *sycp1* mutant females from data in (B). Colored lines represent animals from three broods. (E) Table showing a subset of embryos placed in the nursery from select broods and the percent that survived to adulthood. (F) Examples of normal-looking and deformed embryos from *syce2*^−/−^ mutant mothers. Quantification of the chromosome content in individual normal-looking (MUT-N) and deformed (MUT-D) offspring. The number of chromosomes in 10 nuclei per embryo are shown. Scale bar = 0.5 mm. (G) Examples of normal-looking and deformed embryos from *sycp1*^−/−^ mothers. Quantification of the chromosome content in individual normal-looking (MUT-N) and deformed (MUT-D) offspring. The number of chromosomes in 10 nuclei per embryo are shown. Scale bar = 0.5 mm.

Crosses involving *sycp1^−/−^* males and females also yield similar numbers of spawned eggs (258.0 ± 61.7 and 162.6 ± 87.0, respectively) as wild-type controls (280.1 ± 113.1 and 178.2 ± 181.4, respectively) (Fig 4B). However, fertility is dramatically reduced in *sycp1^−/−^* males (<0.04 % ± 0.1%) compared to controls (75.9 % ± 25%) while the fertility of *sycp1^−/−^* females (82.8 % ± 25.7%) is not significantly different from controls (88.9 % ± 13.5%) (Fig 4B). Over the course of this study, we found that 6 out of 4233 eggs (0.14%) were fertilized by *sycp1^−/−^* males, indicating that completion of meiosis does occur at a low rate. These results reveal sexually dimorphic phenotypes between *sycp1^−/−^* mutant males and females and suggest that the absence of Sycp1 prevents spermatogenesis but not oogenesis.

Remarkably, a high percentage of fertilized eggs from both *syce2^−/−^* males and females appear morphologically healthy at 1- and 5-days post fertilization (dpf) (Fig 4C). In contrast, the majority of offspring from *sycp1^−/−^* mutant females appear malformed at 1 and 5 dpf (Fig 4D). Note, there was some variability between broods and between crosses within the same brood in *syce2^−/−^* mothers which could reflect day-to-day effects on crossing or tank differences between broods. We raised a subset of normal-looking offspring from *syce2^−/−^* and *sycp1^−/−^* mothers past 5 dpf, and a substantial percentage survived into adulthood (Fig 4E). This was surprising given the severity of the synapsis defects, yet it suggested that chromosomes segregate normally at least in a subset of meioses and/or that some degree of gamete aneuploidy is tolerated during development.

To determine whether the mutants produce euploid or aneuploid offspring, we performed metaphase chromosome spreads on 1 day-old normal-looking and malformed offspring from *syce2^−/−^* and *sycp1^−/−^* mothers. Both normal and malformed progeny from *syce2^−/−^* and *sycp1^−/−^* females exhibit considerable variability in the number of chromosomes per nucleus (dots) within individual embryos (bars), while wild-type nuclei rarely deviate from 50 chromosomes (Fig 4F, G). Although the aneuploidy in normal-looking embryos could represent embryos that would have developed abnormally after 1 dpf, approximately half of the embryos from *syce2^−/−^* or *sycp1^−/−^* mothers survive to adulthood (Fig 4E), suggesting that some of the normal-looking embryos develop normally despite having a significant number of aneuploid cells at 1 dpf. The variation in the number of chromosomes in different cells from the same embryo likely reflects somatic mosaicism, where chromosome missegregation occurs after fertilization [60]. These results indicate that defects in the meiotic machinery can lead to genome instability in the developing embryo.

#### Sex ratios are altered in *sycp1^−/−^* mutants but not *syce2^−/−^* and mutants

Sex determination in domesticated zebrafish is poorly understood, but it is not regulated solely by sex chromosomes [61]. Instead, all larvae (~14–20 dpf) regardless of their eventual sex produce early-stage oocytes (Stage IA) and those that produce a threshold number of oocytes continue to develop as females, while those that do not develop as males [62]. In addition, continuous oocyte production is required to maintain the female phenotype after sex determination [63]. Therefore, a male-biased population can be indicative of a defect in oogenesis [56,64–68]. To gauge if sex ratios are altered in *syce2^−/−^* and *sycp1^−/−^* populations we measured the percentage of males in four different broods of each genotype. We found no significant difference in the percentage of males between *syce2^−/−^* mutant (76.0 ± 14.8%) and wild-type (68.3 ± 6.4%) or heterozygous (66.4 ± 6.0%) populations, indicating that sufficient oocytes are present for female sex determination in these three genotypes (Fig 5A).

**Figure 5.**
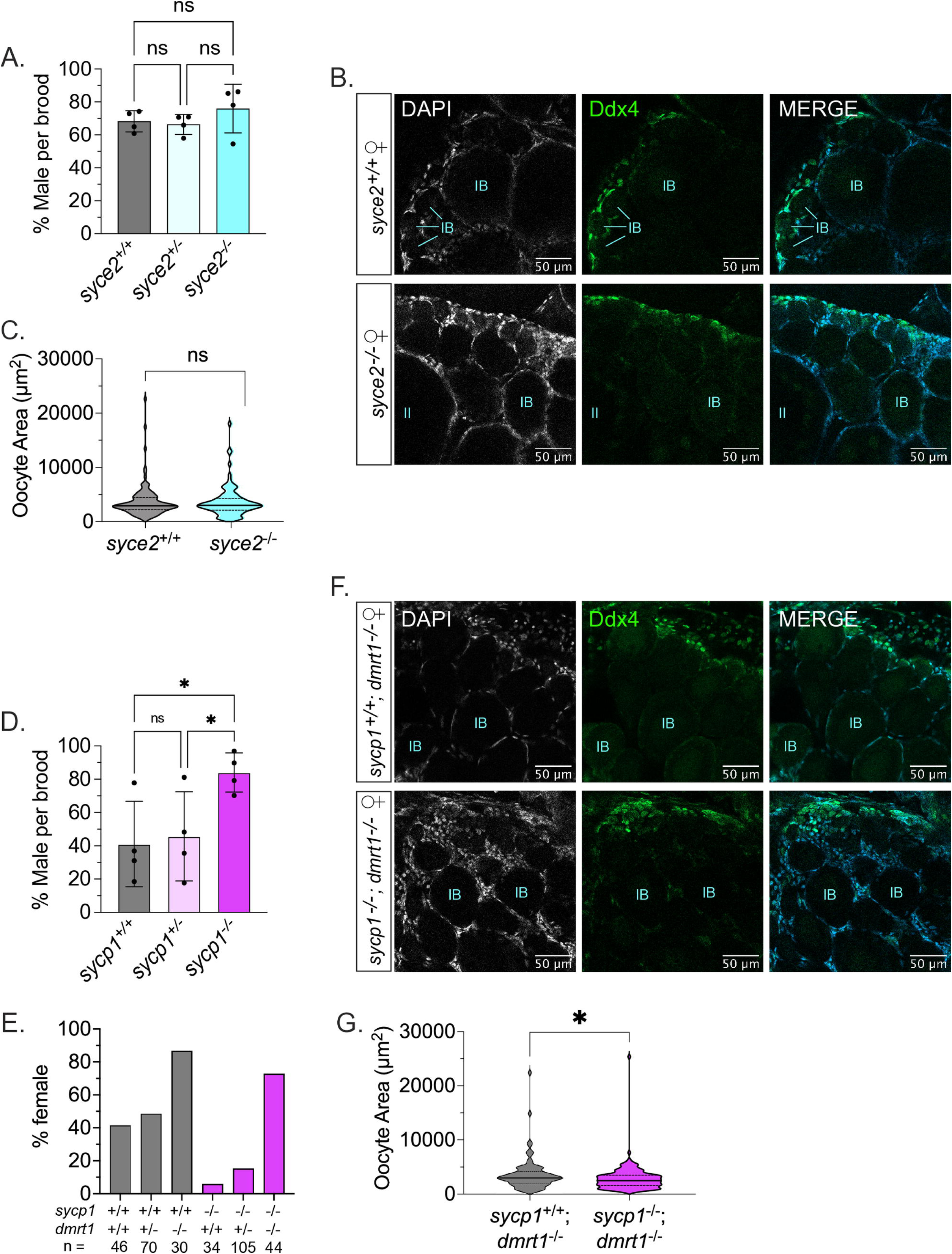
Normal sex ratios and gonad development in *syce2*^−/−^ mutants but not *sycp1^−/−^* mutants. (A) Percentage of animals developing as male in *syce2^+/+^*, *syce2^+/−^* and *syce2*^−/−^ genotypes from four independent experiments (n_1 total_ = 99, n_2 total_ = 342, n_3 total_ = 318, n_4 total_ = 328). Each data point indicates the percentage of males within each genotype per brood. Significance was determined using repeated measures one-way ANOVA testing with Šidák’s multiple comparisons test; ns = not statistically significant. (B) Whole mount ovaries from 45 dpf *syce2*^+/+^ *and syce2*^−/−^ females imaged using confocal microscopy. Ovaries are stained for DAPI (gray) and Ddx4 (green). Scale bar = 50 µm. All images contain the edge of an ovary where cells in early prophase I are found for orientation. (C) Violin plot of the area of individual oocytes from 3 *syce2*^+/+^ and 3 *syce2*^−/−^ whole mount ovaries in (A). n = 118 oocytes (*syce2*^+/+^) and n = 104 oocytes (*syce2*^−/−^). Significance was determined using an unpaired t-test. ns = not significant. (D) Percentage of animals developing as male in *sycp1^+/+^*, *sycp1^+/−^* and *sycp1^−/−^* genotypes from four independent experiments (n_1 total_ = 237, n_2 total_ = 112, n_3 total_ = 167, n_4 total_ = 382). Each data point indicates the percentage of males within each genotype per brood. Significance was determined using repeated measures one-way ANOVA testing with Šidák’s multiple comparisons test; ns = not statistically significant, * = p<0.05. (E) Percent females in *sycp1*^+/+^ and *sycp1*^−/−^ populations that are wild-type, heterozygous or mutant for *dmrt1*. n values for each population are listed in the graph. (F) Whole mount ovaries of *sycp1*^+/+^ and *sycp1*^−/−^ females in a *dmrt1*^−/−^ background stained with DAPI and Ddx4. Scale bar = 50 µm. All images contain the edge of an ovary where cells in early prophase I are found for orientation. (G) Violin plot of the area of individual oocytes from 3 *sycp1*^+/+^; *dmrt1*^−/−^ and 6 *sycp1*^−/−^; *dmrt1*^−/−^ whole mount ovaries in (C). n = 81 oocytes (*sycp1*^+/+^; *dmrt1*^−/−^) and n = 151 oocytes (*sycp1*^−/−^; *dmrt1*^−/−^). Significance was determined using an unpaired t-test. ns = not significant.

To test if oocytes appeared normal, we stained whole gonads from 45 dpf juveniles with an antibody to the germ cell marker Ddx4 and DAPI to visualize DNA. We found *syce2^+/+^* and *syce2^−/−^* ovaries have similar morphologies harboring Stage IB oocytes (Fig 5B), and mutant oocytes are similar in size to wild-type oocytes (Fig 5C) suggesting that aberrant oocytes are not eliminated by apoptosis.

In contrast, the *sycp1^−/−^* mutant population exhibited a significantly higher percentage of males (84.0 ± 11.7%) compared to wild-type (41.1 ± 25.6%) and heterozygous (45.7 ± 26.8%) populations (Fig 5D). These effects show that mutation of *sycp1^−/−^* may be impacting oogenesis, but not to the extent of *sycp1^isa^*^/isa^ where a complete male population was reported [54]. It is possible that differences in animal husbandry and/or strain background (AB, this study, versus India) resulted in insufficient numbers of oocytes to support female sex determination or maintenance in the *scyp1^isa^*^/isa^ mutant.

#### Extending the sex-determination window by mutating *dmrt1* rescues the male bias of *sycp1^−/−^* mutants

We reasoned that if the male bias of *sycp1^−/−^* populations is due to insufficient oocyte production, then expanding the sex-determination window might increase the percentage of females. By contrast, if oocytes are eliminated by checkpoint-mediated apoptosis, then extending the sex determination window will have little or no effect on sex ratios. The sex-determining window depends on the rate of apoptosis of early-stage oocytes, after which point the bipotential juvenile ovary transitions to a testis; mutation of the testis-promoting gene *dmrt1* reduces the number of apoptotic oocytes and skews sex ratios to favor female development [69]. We found that the *dmrt1* mutation dramatically increases the percentage of females in the *sycp1^−/−^* background from 5.9% to 72.7% (Fig 5E). These results suggest that the male bias in *sycp1^−/−^*mutants is due to a delay in reaching the Stage IB stage of oogenesis when female sex is determined and not the result of oocyte elimination by apoptosis stimulated by a strong prophase checkpoint response.

To rule out the possibility that Stage IB oocytes are eliminated from *sycp1^−/−^* ovaries by apoptosis we visually inspected stained whole gonads from 45 dpf juveniles with an antibody to the germ cell marker Ddx4 and DAPI to visualize DNA. Stage IB diplotene oocytes from *sycp1*^−/−^; *dmrt1*^−/−^ gonads were present near the edge of the ovary, where early prophase cells are located, without signs of apoptotic nuclei (Fig 5F). Notably, oocytes are on average smaller than *sycp1*^+/+^; *dmrt1*^−/−^ oocytes (Fig 5G), The smaller oocytes in the *sycp1*^−/−^; *dmrt1*^−/−^ double mutant may be due to a delay in passage through early prophase events that delays the transition from oocytes in a cyst (Stage IA) to those in the follicle (Stage IB) [70]. This delay might explain the male bias of the *sycp1*^−/−^ population. Alternatively, the smaller oocyte size could be due to gonads that start transitioning to testes, but without *dmrt1* revert to an ovary fate leading to female development [69]. An analysis of homolog pairing and synapsis in *sycp1*^−/−^; *dmrt1*^−/−^ surface spread oocytes shows that the chromosome co-alignment phenotype is similar to *sycp1*^−/−^ oocytes (Fig S3; Fig 3D).

### Spermatogenesis is blocked in *sycp1^−/−^* but not *syce2^−/−^* mutants

Next, we examined testes of *syce2^−/−^* and *sycp1*^−/−^ males to test if spermatogenesis is disrupted in these mutants. Consistent with the normal fertility of *syce2^−/−^* males, mutant testes have prominent sperm clusters and are morphologically indistinguishable from wild-type (Fig 6A). By contrast, *sycp1*^−/−^ testes contain no sperm clusters arguing that spermatogenesis is incomplete (Fig 6B). We did, however, observe irregularly condensed nuclei characteristic of metaphase spermatocytes in both wild-type and mutant testes (Leal et al. 2009). To test if these are cells in metaphase, we immunostained wild-type and *sycp1^−/−^* testes with the metaphase marker phospho-Histone H3 (Serine 10) [71] and observed staining of these irregularly shaped nuclei (Fig 6B). This reveals that *sycp1^−/−^* spermatocytes can transit prophase I to reach metaphase and suggests that a prophase I checkpoint leading to arrest is not activated by asynapsis.

**Figure 6.**
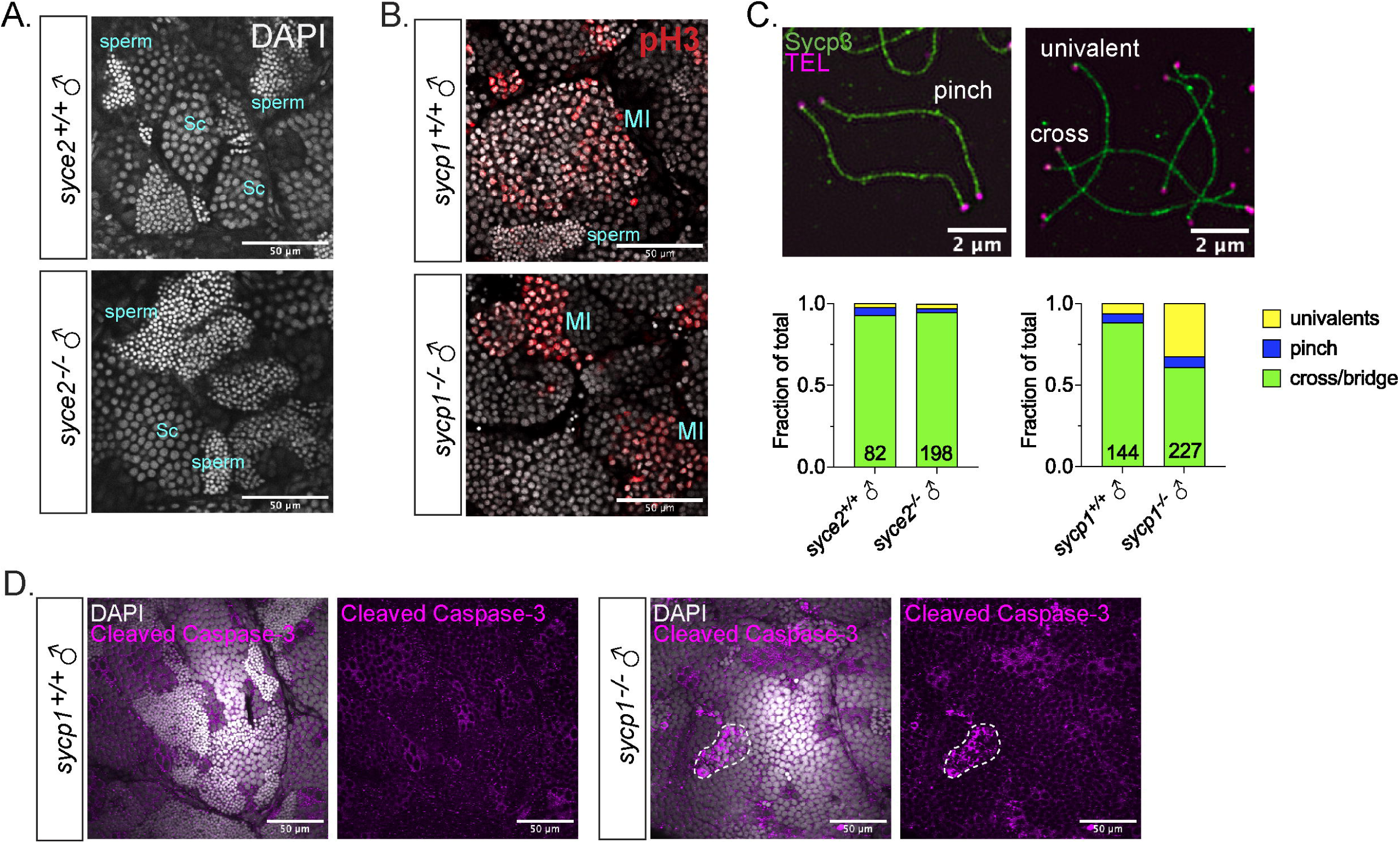
Spermatogenesis arrests at metaphase I in *sycp1^−/−^* males but not *syce2^−/−^* males. (A) Whole mount testes of *syce2*^+/+^ and *syce2*^−/−^ males stained with DAPI. Examples of spermatocytes in meiosis I prophase (Sc) and sperm are noted. Scale bar = 50 µm. (B) Whole mount testes of *sycp1*^+/+^ and *sycp1*^−/−^ males stained with DAPI and phospho-Histone H3 (pH3; red). Examples of spermatocytes in metaphase (MI) and sperm are noted. Scale bar = 50 µm. (C) Proportion of distinguishable chromosome configurations (cross/bridge bivalents, pinch bivalents and univalents) among diplotene (wild type), pachytene-like (*syce2*^−/−^) and pachytene-diplotene-like (*sycp1*^−/−^) chromosomes. n = 82, *syce*^+/+^; n = 198, *syce2*^−/−^; n = 144, *sycp1*^+/+^; n = 227, *sycp1*^−/−^. Data is compiled from seven independent experiments. (D) Whole mount testes of *sycp1*^+/+^ (A) and *sycp1*^−/−^ (B) males stained with DAPI (gray) and Cleaved Caspase-3 (magenta). Dashed lines represent apoptotic nuclei. Scale bar = 50 µm.

### Univalents are more common in *sycp1^−/−^* than *syce2^−/−^* mutants

Next, we investigated whether the different reproductive outcomes in *syce2^−/−^* and *sycp1*^−/−^ mutants could be due to a difference in the number of crossovers. We examined chromosome spreads and identified late prophase *syce2*^−/−^ and *sycp1^−/−^* nuclei and wild-type diplotene nuclei. We observed three types of chromosome configuration (Fig 6C): (1) bivalents in a pinched configuration, where sub-telomeric regions are parallel to each other but with no apparent crossing over of axes; (2) bivalents with crossed or bridged axes, where axes overlap each other or appear bridged by a short stretch of Sycp3; or (3) univalents, where a full-length chromosome is found with no apparent partner. Because of the tangled nature of *syce2^−/−^* and *sycp1*^−/−^ spread chromosomes, we limited our analysis to only those chromosomes where the full axes could be unambiguously traced, which were generally on the outer edges of the spread chromosome region. The proportion of univalents in *syce2^−/−^* and *syce2^+/+^* spermatocytes was comparable (3% vs 2.4%, respectively; Fig 6C). This result suggests that short stretches of synapsed-like regions observed in *syce2* mutants are sufficient to promote the formation of at least one crossover between each pair of homologs. By contrast, the proportion of univalents in *sycp1^−/−^* spermatocytes was higher than in *sycp1^+/+^* (32.6% vs 6.3%; Fig 6C).

In mice, univalents cannot be stably oriented on the meiosis I spindle, which subsequently activates the spindle assembly checkpoint (SAC) and leads to apoptosis [72]. To test if apoptotic cells were present in the *sycp1^−/−^* spermatocytes we immunostained mutant testis to visualize the apoptotic marker, cleaved caspase-3, and found several patches of apoptotic cells (Fig 6D). This indicates that the increased incidence of univalents in *sycp1*^−/−^ males leads to metaphase arrest and apoptosis, likely through SAC activation. However, we cannot rule out the possibility that the chromosomes simply cannot separate at anaphase due to entanglements [73].

### Less efficient DSB repair in *syce2^−/−^* and *sycp1^−/−^* mutants

In mice, mutations in *Syce2 and Sycp1* lead to the accumulation of unrepaired DSBs [23,74]. We investigated whether there is a DSB repair defect in the analogous zebrafish mutants by examining spread spermatocyte chromosomes for the presence of Rad51, an indicator of unrepaired DSBs [75]. Although the pachytene- and diplotene-like stages are indistinguishable in the *sycp1*^−/−^ spermatocytes spreads, we were able to identify diplotene-like spermatocytes in *syce2^−/−^* mutants by finding nuclei in which the axes were full-length but Sycp1 localization in synapsed-like regions is largely absent. In wild-type spermatocytes, Rad51 foci are present in leptotene (65.86 ± 30.6) and zygotene (74.53 ± 25.01) with a sharp reduction in pachytene (39 ± 16.02) and diplotene (16.86 ± 7.77) (Fig 7A, D). Both *syce2^−/−^* and *sycp1*^−/−^ mutants have significantly more Rad51 foci from zygotene (109.3 ± 36.97 and 122.5 ± 32.54, respectively) to late prophase (79.86 ± 40.25, P-like; 65.5 ± 25.08, D-like; and 94.81 ± 34.43, P/D-like, respectively) (Fig 7A–D), indicative of increased DSB formation and/or less efficient DSB repair. The number of foci post-zygotene is considerably decreases in both mutants, suggesting that a fraction of DSBs are repaired (Fig 7D). Despite the synapsed-like nature of *syce2^−/−^* homologs, there is no significant difference in Rad51 foci between *syce2^−/−^* and *sycp1*^−/−^ mutants. We then immunostained whole *sycp1^−/−^* mutant testes for the DSB marker γH2AX, which allowed us to assess whether DSBs are still present in metaphase cells. γH2AX is not detectable from metaphase nuclei, suggesting that despite their persistence through much of prophase I, DSBs are eventually repaired (Fig S4A). These data show that both *syce2^−/−^* and *sycp1*^−/−^ mutant males are less efficient at repairing DSBs during prophase. We also immunostained whole juvenile ovaries from *sycp1^−/−^* mutant females for γH2AX. While γH2AX is observed in Stage IA oocytes that are in leptotene/zygotene, γH2AX is not detectable in Stage IB oocytes that reach diplotene (Fig S4B). This indicates that DSBs are repaired by late prophase I in oocytes.

**Figure 7.**
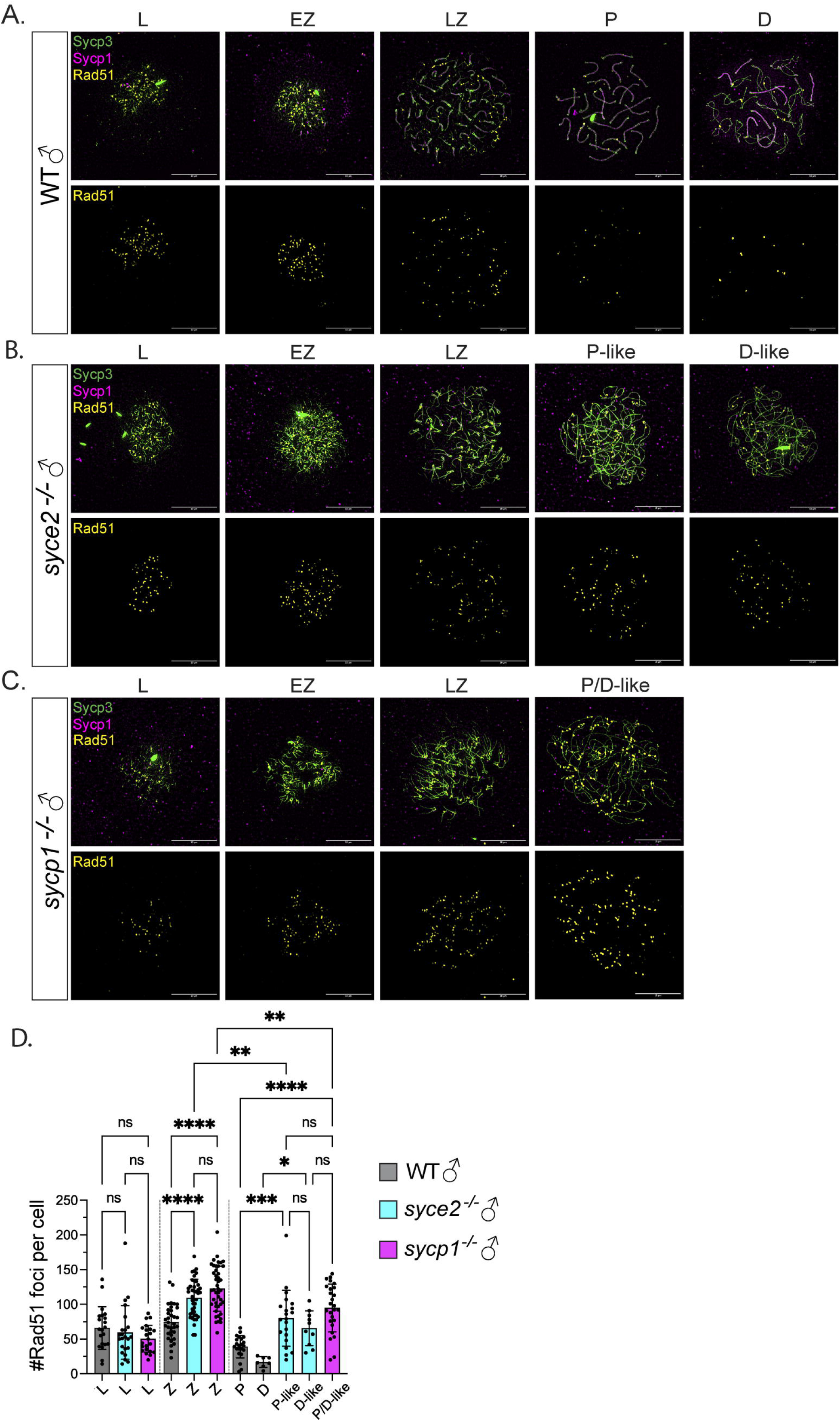
Inefficient DSB repair in *syce2*^−/−^ and *sycp1^−/−^* mutants during prophase. (A–C) Surface-spread chromosomes from wild-type (A), *sycpe2^−/−^* (B) and *sycp1*^−/−^ (C) spermatocytes immunostained for Sycp3 (green), Sycp1 (magenta) and Rad51 (yellow). Examples of spread chromosomes from meiotic prophase: L, EZ, LZ, P, D, P-like (*syce2*^−/−^) and P/D-like(*sycp1*^−/−^). D-like nuclei (*syce2*^−/−^) are categorized as having full-length axes, but Sycp1 is largely absent from pseudo-synapsed regions throughout the nucleus. Scale bar = 10 µm. (D) Number of Rad51 foci per cell using data from (A–C). EZ to LZ nuclei are grouped together as zygotene (Z). Data is pooled from 2 independent experiments. n = 21, WT – L; n = 23, *syce2*^−/−^ – L; n = 22, *sycp1*^−/−^ – L; n = 34, WT – Z; n = 41, *syce2*^−/−^ – Z; n = 44, *sycp1*^−/−^ – Z; n = 22, WT – P; n = 7, WT – D; n = 21, *syce2*^−/−^ – P-like; n = 10, *syce2*^−/−^ – D-like; n = 26, *sycp1*^−/−^ – P/D-like. Significance was determined using ordinary one-way ANOVA testing with Šidák’s multiple comparisons test. ns = not significant; ** = p < 0.01; *** = p <0.001; **** = p < 0.0001.

### Partial synapsis in *syce2^−/−^* mutants is insufficient for Hormad1 removal

In mice, DSB formation is promoted by HORMAD1, an axis-associated HORMA domain containing protein that is removed once a region has become synapsed [76]. In wild-type zebrafish, Hormad1 localization to meiotic chromosomes is transient since SC elongation immediately follows axis formation. Consequently, only short stretches of Hormad1 staining are seen along asynapsed axes in spermatocytes during leptotene and zygotene [54] (Fig 8A). Hormad1 also binds to diplotene chromosomes once the synaptonemal complex is disassembled, similar to what has been observed in mice [77,78], In *syce2*^−/−^ leptotene spermatocytes, Hormad1 localizes to asynapsed axes as seen in mouse (Fig 8B) [23]. Hormad1 persists in axial regions of partially synapsed regions in zygotene and pachytene-like *syce2*^−/−^ spermatocytes resulting in Hormad1 coating entire chromosome axes. Since Sycp1 fluorescence intensity in *syce2*^−/−^ mutants never reaches the same level as seen in wild-type (Fig 2D), it is possible that there is insufficient Sycp1 to displace Hormad1 from partially synapsed regions. However, as shown in a mouse *Trip13* mutant, axis-association of HORMAD1 is not mutually exclusive with the presence of SC [78]. We also found Hormad1 localization along axes in *sycp1*^−/−^ spermatocytes (Fig 8C), as was described previously for the *sycp1*^isa/isa^ mutant [54]. These data are summarized in Fig S5. In mice, Hormad1 staining persists at centromeres of pachytene chromosomes, however, we saw no evidence of this localization on pachytene chromosomes in zebrafish spermatocytes.

**Figure 8.**
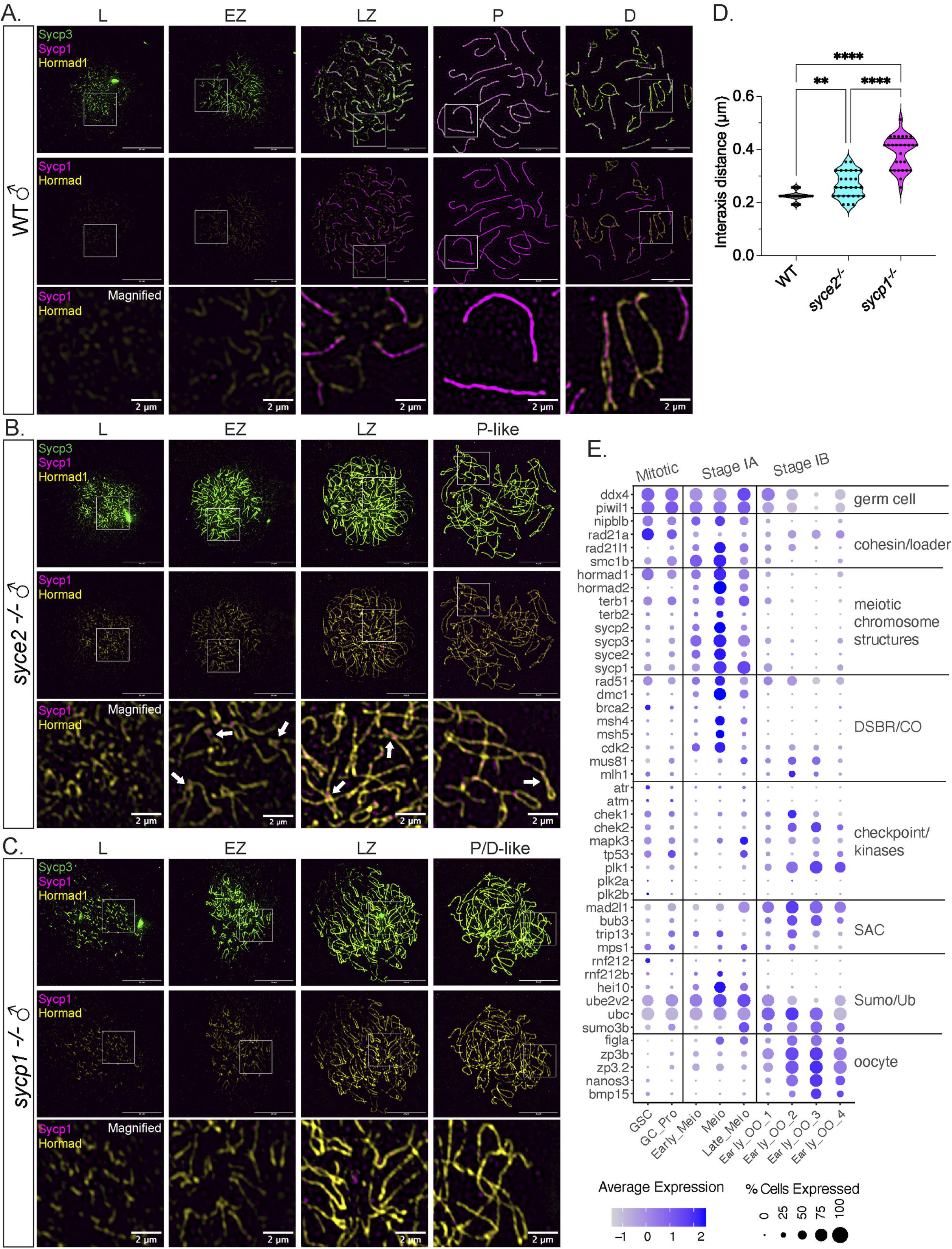
Sycp1 localization to synapsed-like regions is not sufficient for Hormad1 removal. (A–C) Surface-spread chromosomes from wild-type (A), *sycpe2^−/−^* (B) and *sycp1*^−/−^ spermatocytes (C) immunostained for Sycp3 (green), Sycp1 (magenta) and Hormad1 (yellow). Examples of spread chromosomes from meiotic prophase described in Fig 7 A–C. Scale bar = 10 µm. Boxes represent magnified areas for each Stage. Scale bar for magnified examples = 2 µm. Arrows represent Hormad1 localization to pseudo-synapsed regions with Sycp1. (D) Violin plot showing the distance between axes for co-aligned axial pairs in early zygotene for wild-type (n = 28), *syce2*^−/−^ (n = 28) and *sycp1*^−/−^ (n = 28) spermatocytes. 4 cells from each genotype were used for the analysis. Significance was determined using ordinary one-way ANOVA testing with uncorrected Fisher’s LSD test. (E) ScRNA-seq expression of select genes from [102]. Germline stem cell (GSC), oocyte progeniture (OO_Pro), meiotic prophase stage (meio), oocyte (OO).

Given the similarities in Hormad1 localization, DSB repair inefficiency and sub-telomeric co-alignment in *syce2^−/−^* and *sycp1*^−/−^ mutants, we investigated whether the inter-axis distance in the sub-telomeric region is also similar. Since the distance between axes can vary in spreads done on different days, we prepared a set of chromosome spreads from wild-type, *syce2*^−/−^ and *sycp1*^−/−^ spermatocytes in parallel. We focused our analysis on zygotene when chromosome entanglements are infrequent and found that the inter-axis distance in *syce2^−/−^* spermatocytes (0.266 ± 0.049) is intermediate between wild-type (0.224 ± 0.019) and *sycp1^−/−^* (0.387 ± 0.062) (Fig 8D). These results suggest that Sycp1 contributes to close alignment of homologs even in the absence of a key central element protein.

## Discussion

Our study supports four main conclusions regarding meiosis in zebrafish. First, partial synapsis, limited primarily to the ends of chromosome, is sufficient to create fertile gametes that give rise to normal progeny. Second, defects in synapsis do not elicit a strong prophase I checkpoint response in spermatocytes and oocytes. Third, the rate of progression through prophase can influence determination and maintenance of the female sexual fate. Fourth, defects in meiotic prophase I can promote genome instability in developing F1 progeny.

We found that *syce2^−/−^* and *sycp1^−/−^* females produce normal numbers of fertile eggs and progeny that live to adulthood despite extensive asynapsis and inefficient DSB repair during meiotic prophase I. This is in striking contrast to what is seen in mice, where *syce2^−/−^* and *sycp1^−/−^* females are infertile and fail to produce oocytes that progress past the pachytene stage [23]. *syce2^−/−^* zebrafish males are also fertile and produce largely normal progeny. This is similar to yeast central element mutant *ecm11Δ* that produces viable spores despite asynapsed homologs, presumably because the ZMM pathway is intact and obligate crossovers are executed [28]. In contrast *sycp1^−/−^* spermatocytes transit prophase and do not arrest until metaphase I. The presence of *sycp1^−/−^* apoptotic cells at this stage is presumably caused by univalents that activate the spindle-assembly checkpoint (SAC). Therefore, our study provides evidence that asynapsed chromosomes fail to elicit a strong prophase I checkpoint operating in oocytes or spermatocytes, making zebrafish more similar to yeast [39–41] or *Drosophila* females [44,45] compared to mice and *C. elegans* where asynapsis leads to prophase I arrest and apoptosis [31–37].

Why might zebrafish have a weak meiotic prophase I checkpoint compared to mammals? One distinguishing feature of mice and *C. elegans* compared to other models is the meiotic silencing of unpaired chromatin (MSUC) response, which is related to meiotic sex chromosome inactivation (MSCI) that silences unpaired regions of sex chromosomes to evade checkpoint activation [79,80]. Self-pairing of folded supernumerary B chromosomes and nonhomologous synapsis of sex chromosomes may play a role in circumventing a MSUC activation in fish [81–83]. Alternatively, a MSUC pathway may not exist. Evidence of both male and female heterogamety (e.g. XY and ZW) implies switching from one system to another (e.g. genus *Oryzias*), which might be enabled by a weak synapsis checkpoint [84].

Weak checkpoints in the teleost lineage could be a vestige of the whole genome duplication event that occurred 320-350 million years ago giving rise to a common polyploid ancestor [85]. Moreover, more recent genome doubling (autoploidy) and hybridization between different species (allopolyploidy) is common among fish species [86]. Polyploidy could create challenges for meiotic homolog pairing and recombination and these challenges might be better tolerated with a relaxed checkpoint system [87]. Polyploidy and whole genome duplication have occurred multiple times in plants, often contributing to speciation [88]. The relaxed nature of meiotic checkpoints in plants is often attributed to the extensive polyploidization across plant species [87]. *Arabidopsis* mutants with asynapsis and unrepaired DSBs are still capable of completing meiosis, suggesting a weak checkpoint [89]. Though, at elevated temperatures a specialized variation of the prophase I checkpoint appears to exist [90]. Among other vertebrate species, polyploidy is most common in amphibians and fishes, [91] but no undisputed evidence of a mammalian polyploid species exists [92]. It has also been suggested that reduced checkpoint stringency could favor animal taxa that produce large numbers of progeny in fresh water conditions where fertilization occurs outside of the body [93,94].

The ability to compare males and females allowed us to detect sex-specific differences in chromosome dynamics, checkpoint response and reproduction. We showed that extending the window of sex determination, by inhibiting male differentiation, results in a dramatic rescue of the male-skewed *sycp1* population. This suggests the male skew may not be due to an activation of a canonical meiotic checkpoint that would result in oocyte apoptosis and sex reversal, but instead oocyte numbers simply fail to reach a threshold for female development before the close of the sex-determination window in a subset of animals [62]. Indeed, the smaller oocytes seen in *sycp1^−/−^*; *dmrt1^−/−^* double mutants compared to *dmrt1^−/−^* single mutants may indicate that progression through early meiotic prophase I (e.g. leptotene through pachytene) is slower in the absence of *sycp1*, perhaps due to slower pairing kinetics, DSB repair, removal of entanglements, or by transient recombination checkpoint activation. The previously described *sycp1^isa/isa^* mutant develops into an entirely male mutant population with evidence of female-to-male sex reversal [54]. We attribute the differences between alleles to animal husbandry or strain background (AB versus India) where the threshold number of oocytes for female fate determination in the *scyp1^isa/isa^* mutant background might be higher. We predict that mutating *dmrt1* and prolonging the sex determination window would also suppress female-to-male sex reversal phenotype of *sycp1^isa/isa^* mutants.

Female-to-male sex reversal has been observed in several zebrafish mutants with defects in meiotic processes including, the chromosome axis genes s*ycp2* [66] and *sycp3* [95], the meiosis-specific cohesin subunits *smc1β* [96] and *rad21l1*[67], and the DSB repair genes *brca2* [65,68], *rad51* [97] and *fancl* [64] and other Fanconi anemia genes [56] affecting a range of phenotypes from shortened axial structures, reduced DSB formation, unrepaired DSBs and asynapsis. The lack of females is partially rescued in *sycp3*, *rad21l1*, *brca2*, *rad51* and *fancl* mutants by re*ducing* Tp53-mediated apoptosis [64,65,67,68,95,97]. Since oocyte production is critical to female sex determination, it was reasoned that inhibiting apoptosis could increase the threshold number of oocytes for the female developmental pathway [64]. In mice, the Tp53 ortholog is involved in eliminating cells with unrepaired DSBs [98], so it appears likely that Tp53 in zebrafish is promoting apoptosis due to the presence of unrepaired DSBs in these mutants [47,99]. Since *sycp1^−/−^; dmrt1^−/−^* juveniles achieve female-sex fate with a Tp53-mediated checkpoint pathway presumably intact, we argue that any remaining DSBs in the *syce2* or *sycp1* mutants at the time of sex determination are repaired or are not present at sufficient levels to stimulate a strong checkpoint arrest [69].

In mice, CHEK1 and CHEK2 checkpoint kinases play semi-redundant roles in meiosis by identifying and eliminating oocytes with unrepaired DSBs by signaling to their phosphorylation targets TRP53 and TAp63 [100]. Similarly, RAD53, a CHEK2 homolog in budding yeast, mediates repair of DSBs that have escaped the meiotic recombination checkpoint [101]. Interestingly, published scRNA-seq data show that zebrafish *chek1* and *chek2* are expressed at *low* levels in the cell cluster where chromosome axis and SC genes are also expressed yet they are expressed at relatively *high* levels in the cell cluster containing transcripts characteristic of dictyate-arrested cells such as *figla*, *gdf9*, and *bmp15*, *zp3b, zp3.2*, and *nanos3* [99,102]. Expression of the DSB repair gene *rad51* also persists in cells expressing *chek1* and *chek2* suggesting these genes might play a role signaling the presence of unrepaired DSBs to a Tp53-mediated DNA damage response. In yeast, *C. elegans*, and mice, DSB repair in meiosis prophase I can also be divided into two stages with the first biasing repair of DSBs with the homologous chromosome and a second stage changing the bias to Rad51-mediated sister-chromatid directed repair to repair remaining DSBs [42,103–107]. It is possible that these two stages could be demarcated as Stage IA and Stage IB oocytes. Interestingly, Stage IB oocytes can be found in the *scyp1^isa/isa^* juveniles that will eventually undergo sex reversal [54]. Presumably these animals make sufficient oocytes for female-fate determination. Thus, the *sycp1^isa/isa^* mutant may make enough oocytes for female fate determination but not enough for female maintenance, possibly due to an active “second phase”-like *Chek2/Tp53* mediated apoptosis in diplotene cells. These data do not preclude the presence of a first-stage meiosis I DSB checkpoint since DSB repair mutants *brca2* develop solely as males with arrested spermatocytes yet do not appear to form Stage IB oocytes in the juvenile ovary [65].

Chek2/Chk2/Mek1 orthologs play a key function as a meiosis prophase I checkpoint kinase in mouse, *C. elegans*, *Drosophila* and budding yeast and mediate checkpoint-induced arrest in yeast, and arrest and apoptosis in *C. elegans* and mice with synapsis defects [34,41,100,108–111]. Thus, the apparent absence of a synapsis checkpoint in zebrafish is consistent with low expression of *chek2* during early meiotic prophase. In *C. elegans* the synapsis checkpoint operates by the activation of the CHK-2 kinase by HORMA domain proteins found on unsynapsed axes [112]. That there is abundant Hormad1 protein on both partially synapsed and asynapsed chromosomal regions in the *syce2^−/−^* spermatocytes, yet they progress to form fertile sperm, supports the notion that Hormad1 function in zebrafish can be uncoupled from a prophase checkpoint function.

Analysis of germline-expressed genes in the ovary [102] points to a genetically modulated transition that occurs between Stage IA oocytes in early prophase (leptotene-pachytene) and the Stage IB oocytes in dictyate arrest (Fig 8E). As mentioned above, *chek1/chek2* are not expressed in early prophase but are elevated in the Stage IB oocytes reaching dictyate arrest. Meiotic transitions in *C. elegans* are regulated by the polo-like kinases PLK-1 and PLK-2 and the extracellular signal– regulated kinase ERK [113–115]. Zebrafish homologs *plk1, plk2* and the ERK homolog, *mapk3,* show varying degrees of expression; *plk1* is found at high levels in Stage IB cells, but *plk2a* and *plk2b* are only present in a small number of cells and at low levels. Interestingly, *mapk3* appears to be present in the Stage IA population, so this kinase could potentially modulate the transition to Stage IB. SUMO- and ubiquitin-mediated regulation of crossover control during pachytene in mice is mediated by SUMO E3 ligases RNF212 and RNF212b and the ubiquitin E3 ligase HEI10 [116–119]. Expression of the zebrafish *HEI10* homolog *ccnb1lp1* is enriched in Stage IA oocytes that also express the crossover assurance genes *msh4, msh5*, and *cdk2*. *rnf212*b but not *rnf212* is expressed at high levels in a fraction of Stage IA oocytes.

While a prophase I checkpoint may not be stringent in zebrafish, the accumulation of metaphase I spermatocytes in *sycp1* mutant males suggests that there is a robust SAC in this sex. In *cntd1* and *mlh1* mutant males, spermatocytes reach the pachytene stage with fully synapsed chromosomes but are defective in forming crossovers. In these cases, spermatocytes arrest at metaphase I, triggering apoptosis [120,121]. Other evidence for the SAC comes from studies showing metaphase arrest and apoptosis in testes treated with nocodazole at an elevated temperature; in this case, apoptosis is suppressed by mutation of the checkpoint kinase gene *mps1* [122]. Both male and female zebrafish homozygous for the hypomorphic *mps1* allele display germ cell aneuploidy suggesting that *mps1* may play other roles in limiting segregation errors during germ-cell development. By contrast, the SAC does not seem to be activated in *syce2^−/−^* males, consistent with the low numbers of univalents we observed in chromosome spreads.

Though the SAC operates in human and mouse oocytes [123,124] – albeit in a less permissive state compared to spermatocytes [125] – we did not see diminished numbers of unfertilized eggs from *scyp1^−/−^* females. One possible reason to explain a weak or absent SAC could be the large oocyte volume compared to mouse and human oocytes. It has been suggested that oocyte size can lower the accuracy of the SAC by lowering the concentration of SAC proteins and weakening the checkpoint [126]. Indeed, Xenopus oocytes lack the SAC altogether [127]. In zebrafish, oocytes grow to 690 µm before metaphase I compared to 80-150 µm for mammalian oocytes [70], which could account for dilution or sequestration of SAC factors. Zebrafish homologs of genes associated with the meiotic SAC including *mad2l1*, *bub3*, and *trip13* [125,128] are found expressed in Stage IB oocytes (Fig 8E).

Finally, one aspect of our work that was surprising was the high degree of somatic mosaic aneuploidy observed among cells of individual embryos of *syce2^−/−^* and *sycp1^−/−^* mothers. In humans, mosaic aneuploidy occurs in at least 80% of human blastocyst-stage embryos, often with less than 20% of cells showing defects [60]. The apparent lack of prophase I checkpoints in zebrafish make this an excellent model to explore how defects in meiosis contribute to this phenomenon.

## Materials and methods

### Ethics Statement

The UC Davis Institutional Animal Care and Use Committee (IACUC) has approved of this work under the protocol #23636; For noninvasive procedures (e.g. fin clips for genotyping), zebrafish were anesthetized using tricaine. Invasive surgical methods were performed on fish euthanized by submerging fish in ice water.

### Zebrafish husbandry and strains

Zebrafish husbandry was performed as previously described [102] with the following modifications. At 5 days post fertilization (dpf), zebrafish larvae were placed into 1 L aquaria at 40 fish/tank. At 5–10 dpf larvae were fed ~3 mL rotifers *ad libitum*. From 10– 14 dpf larvae were fed 3 mL rotifers *ad libitum* and 1 drop concentrated brine shrimp twice daily. From 14–23 dpf, larvae were fed 2 drops concentrated brine shrimp twice daily. From 23–30 dpf, larvae were fed 3 drops concentrated brine shrimp twice daily. At 30 dpf, juvenile fish were moved to 9 L tanks continuous system water flow and no more than 40 fish. From 30–60 dpf, juveniles were fed ~500 µL concentrated brine shrimp and 100 mg Gemma Micro 300 twice daily. Starting at 60 dpf, fish were fed 100 mg Zebrafish Select Diet twice daily.

### Mutant generation and identification

The wild-type AB strain was used to generate the *syce2^−/−^* and *sycp1^−/−^* mutants. All fish used in the experiments were outcrossed to the AB strain. The *syce2* and *sycp1* mutants were generated with CRISPR/Cas9 using the CRISPR Design Tool (Synthego, Redwood City, CA, USA). gRNA target sequences: 5’ UUCACCAGCCAACAAUACAG 3’ (*syce2*, exon 4) and 5’ CGAGCAGUUUGGAGUACAGC 3’ (*sycp1*, exon 5). One-cell embryos were injected with 300 ng/µL gRNA, 2µM Cas9-NLS purified protein (MacroLab, University of California, Berkeley), and phenol red (5% in 2M KCl). Injected founder fish were raised to adulthood and outcrossed to wild-type fish. Genomic DNA was extracted from the caudal fin of resulting offspring to screen for mutations via High Resolution Melt Analysis (HRMA) (Blokhina et al. 2019). *syce2* HRMA primers: forward 5’-TGATGATTCAGGAATTGGTGTT-3’; reverse 5’-CATCTATTGGTGAATTGTTTGGAG-3’. *sycp1* HRMA primers: forward 5’-GGAGCTTTTTGTTGTTGTTGCAT-3’; reverse 5’-ATCGGTTCTGAACTTCCAGAGT 3’. Mutations were identified in offspring of *syce2* injected fish using PolyPeak Parser (Hill et al. 2014) after Sanger Sequencing of PCR products amplified by Phusion DNA polymerase (New England Biolabs catalog #M0530L). Mutations were confirmed by sequencing PCR products from homozygous mutants. PCR primers: forward 5’-ACCATTTCGTTTCTTGTGTCAGT-3’; reverse 5’-ACACGTTTTACTAATCTGCCTGT-3’. Mutations were identified in offspring of *sycp1* injected fish using PCR products amplified by Phusion DNA polymerase and cloned into the pCR Blunt II-TOPO vector (Invitrogen catalog #450245). *sycp1* PCR primers: forward 5’-GCCAATGGAAAAGGAAGAGGTC-3’; reverse 5’-TGCTTAGACTTTCATTTGCGAACT-3’. The *dmrt1* mutation was generated as previously described (Webster et al. 2017).

### Genotyping

*syce2* and *sycp1* fish were genotyped using the HRMA primers above. *dmrt1* HRMA primers (BWD934 and BWD935) are reported elsewhere [69] (Webster et al. 2017).

### Protein alignment

Protein alignments were done using the Clustal Omega multiple sequence alignment tool [129] using protein sequences from the UniProtKB database for zebrafish, human, and mouse Syce2 orthologs (Q56P19, Q6PIF2 and Q505B8, respectively) and Sycp1 orthologs (F1QYS1, Q15431 and Q62209, respectively).

### RT-PCR

Total RNA extraction from individual testes was performed as previously described (Blokhina et al. 2021). Total RNA extraction from individual testes was performed as previously described with the following modification: upon liquid phase separation 70 µL of the top, clear aqueous solution was purified using the RNA Clean and Concentrator-5 Kit (Zymo Research catalog #R1015). *syce2* RT-PCR primers: forward 5’-CATACATCAGCAAAGGTGCCAG-3’; reverse 5’-CACCAAAAGGTTCCAGGGCT-3’. *sycp1* RT-PCR primers: forward 5’-CGTCGAGAAGCTTTGGTGACT-3’; 5’-TGCCACCATCCTCTGAACAT-3’. *eef1a1l1* (control) RT-PCR primers: forward 5’-CTACCTACCCTCCTCTTGGTCG-3’; reverse 5’-CCTTAAGTAGAGTGCCCAGGT-3’.

### Sex ratios

The sex of adult zebrafish (> 60 dpf) was determined by the absence or presence of an ovipositor found only in females. For juvenile animals between ~45–52 dpf, the sex was determined by the presence of Stage IB oocytes in the follicle stage.

### Chromosome spread preparation and staining

Four to seven testes from adult males (> 60 dpf) were dissected and prepared as previously described (Blokhina et al. 2019) with minor modifications in reagents used: Trypsin inhibitor (Sigma-Aldrich, catalog #T9253); Paraformaldehyde (ThermoFisher, catalog # 28908). Five to fifteen ovaries were dissected from juvenile females (46–52 dpf) and prepared as previously described (Blokhina et al. 2019).

PNA telomere probe and primary and secondary antibody staining was performed as previously described (Blokhina et al. 2019). PNA Telomere probe used: TelC-Alexa647 (PNA Bio Inc, catalog #F1013). Primary antibodies used: 1:200 rabbit anti-Sycp3 (Abcam, catalog #ab150292, discontinued); 1:200 rabbit anti-Sycp3 (Novus Biologicals, catalog #NB300232); 1:100 chicken anti-Sycp1 (Blokhina et al. 2019); 1:100 mouse anti-Rad51 (ThermoFisher catalog# MA5-14419, discontinued); 1:100 mouse anti-Hormad1 (Imai et al. 2021). Secondary antibodies used at 1:1000: goat anti-rabbit IgG Alexa Fluor 488 (ThermoFisher, catalog #A-11008); goat anti-chicken IgY Alexa Fluor 568 (ThermoFisher, catalog #A-11041); goat anti-chicken IgY Alexa Fluor 594 (ThermoFisher, catalog #A-11042); goat anti-mouse IgG Alexa Fluor 647 (ThermoFisher, catalog #A-21236). Slides not stained for telomeres were rehydrated in ~500 µL 1X PBS and then proceeded to primary antibody staining. Slides were mounted with Prolong diamond antifade mountant with DAPI (ThermoFisher, catalog #P36966), Prolong diamond antifade mountant (ThermoFisher, catalog #P36970) or Prolong glass antifade mountant with NucBlue (ThermoFisher, catalog #P36985).

### Whole mount gonad and staining

Adult testes (> 60 dpf) were dissected and stained as previously described (Blokhina et al. 2019). Juvenile ovaries (39-52 dpf) were dissected and stained similarly to adult testes. Primary antibodies used: 1:2000 chicken anti-Ddx4 (Blokhina et al. 2019); 1:200 mouse anti-pH3 (Abcam, catalog #ab14955); 1:100 rabbit anti-γH2AX (Rhodes et al. 2009); 1:200 rabbit anti-γH2AX (GeneTex, GTX127342) rabbit anti-Cleaved-Caspase-3 (Asp175) (Cell Signaling Technology, catalog #9661S). Secondary antibodies used at 1:300: goat anti-chicken IgY Alexa Fluor 488 (ThermoFisher, catalog #A-11039); goat anti-mouse IgG Alexa Fluor 594 (ThermoFisher, catalog #A-11032); goat anti-rabbit IgG Alexa Fluor 594 (ThermoFisher, catalog #A-11012).

### Metaphase spreads

Embryos at ~19–32 hours post fertilization were manually dechorionated and prepared according to [130] with the following modifications: Embryos were incubated in colchicine for 2 hours 15 minutes. The yolk was removed within 16 minutes followed by incubating on ice for 16 minutes. Embryos were mechanically homogenized for 3 minutes in 50 μL of fixative solution followed by brief centrifugation for 1 minute. Instead of pre-equilibrating slides to 4 °C, slides were held face down immediately above a 55°C water bath for 15–20 seconds. The entire cell suspension from one embryo was then dropped onto the center of the slide held at a 30–45° angle to allow the cell suspension to spread. The slide was held face down above a 55°C water bath for 15–20 seconds, air dried for 2 minutes, passed back and forth over a flame 5 times and washed in a coplin jar with 1X PBS for 5 minutes. Slides were air dried or excess PBS was removed with a nasal aspirator to facilitate drying. Slides were stained with Vectashield antifade mounting medium with DAPI (Vector Laboratories, catalog #H-1200) and covered with a glass coverslip.

### Imaging

Images of stained meiotic chromosomes and gonads were collected at the Department of Molecular and Cellular Biology Light Microscopy Imaging Facility at UC Davis. Chromosomes spreads were imaged using the Nikon N-SIM Super-Resolution microscope in 3D-SIM imaging mode with APO TIRF 100X oil lens. Images were collected and reconstructed using the NIS-Elements Imaging Software. Gonads were imaged using the Zeiss 980 Laser Scanning Microscope with Airyscan using LMS confocal. Gonads immunostained for γH2AX and cleaved Caspase-3 were imaged using the Olympus FV1000 laser scanning confocal microscope. All images were processed using Fiji ImageJ software. Only linear modifications to brightness and contrast of whole images were applied. Images of metaphase chromosomes were collected on a Zeiss MetaSystems Imager Z2 microscope 63X lens with MetaFer 5.

### Quantification of mean synapsed length and Sycp1 fluorescence intensity

Mean-synapsed lengths per cell were determined by tracing stretches of Sycp1 lines using the segmented line tool in ImageJ. Since Sycp1 staining is often discontinuous between co-aligned axes in *syce2* mutants, we included regions where co-aligned axes were in a partially synapsed-like configuration (i.e. separated by ~0.2 µm). To calculate the mean fluorescence intensity of Sycp1 signal per cell, we subtracted a mean fluorescence intensity of six boxes taken from the background region of each spread from the mean fluorescence intensity of each scanned Sycp1 segment from the same spread. The average of the adjusted means for each nucleus was normalized to the value calculated from the wild-type EZ spreads to get the relative levels for mutant and wild-type at each stage.

### Interaxis distance

The interaxis distance was measured by drawing a line perpendicular to parallel co-aligned axes in ImageJ to generate kymographs. We identified the maximum intensity signals from the kymographs to determine x-values of the two highest peaks. The x-values were subtracted to calculate the interaxis distance.

### Number of co-aligned sub-telomeric regions

Axes were considered co-aligned if parallel regions of the axes were separated by less than 0.5 µm. In wild-type spermatocytes, synapsed regions were included as co-aligned. Slightly misaligned axes in otherwise parallel axes were considered as co-aligned since the orientation of the chromosomes may shift when the 3D nucleus is flattened to 2D. Axes that were perpendicular to each other but had no other parallel region between them were not considered co-aligned. To calculate the proportion of co-aligned sub-telomeric regions in wild-type diplotene spermatocytes, we analyzed bivalents with asynapsed sub-telomeric regions. Since there is no synapsis in *sycp1* mutants, all possible pairs of co-aligned sub-telomeric regions were included in our analysis.

### Test crosses and embryo tracking

To analyze fertility, individual mutant fish were placed in a divided mating tank overnight with one or two AB wild-type fish of the opposite sex. The divider was removed soon after the onset of light, and any eggs produced were collected with a strainer and categorized as fertilized, unfertilized or decomposed. Fertilized eggs were placed in a petri dish at 28–30 °C with embryo media and monitored at 1 and 5 dpf for malformities.

### Oocyte area

The segmented line tool of ImageJ was used to trace an outline of individual diplotene oocytes in single optical sections in intact gonads imaged using the Zeiss 980 Laser Scanning Microscope. Ovaries were imaged with one edge in the field of view for comparison between images. The area is determined by the number of pixels bound by the perimeter and the pixel density of the image.

### Statistics

All statistical analyses were performed using GraphPad Prism 10. Statistical tests are reported in the figure legends. Comparisons between controls and mutants for individual experiments were done using fish born on the same day.

## Acknowledgements

We thank Kelly Komachi, Nancy Hollingsworth and Satoshi Namekawa for discussions and comments on the manuscript. The Hormad1 antibody was a gift from Yukiko Imai and Norioshi Sakai from the National Institute of Genetics, Mishima Japan.

---

**Supplementary Figure S1.** Targeted mutation of *syce2* and *sycp1* via CRISPR/Cas9.

(A) Creation of the *syce2* mutation by CRISPR/Cas9 showing a 2 nucleotide (nt) deletion within exon 4 (out of 5). The CRISPR target site (green) and premature stop codon (yellow) are shown.

(B) Creation of the *sycp1* mutation by CRISPR/Cas9 showing the site of the mutation within exon 5 (out of 32). The CRISPR target site (green), premature stop codon (yellow) and the complex mutation (orange) are shown. The complex mutation leads to a net increase of 22 nt.

(C–D) RT-PCR from wild-type, *syce2*^−/−^ and *sycp1*^−/−^ testes. Two biological samples were used for each genotype. -RT = no reverse transcriptase. *eef1a1l1* is the positive control.

(D) Note the slower migration in the *sycp1*^−/−^ samples due to the complex mutation with a net increase of 22 nt.

**Supplementary Figure S2.** Identification of wild-type diplotene spermatocytes.

Examples of diplotene spermatocytes from *sycp1*^+/+^ males stained for Sycp3 (green), Sycp1 (magenta and gray) and Telomeres (cyan). Diplotene spermatocytes have full-length chromosome axes with large scale asynapsis. Scale bar = 10 µm.

**Supplementary Figure S3.** *dmrt1* mutation does not affect chromosome co-alignment.

(A–B) Surface-spread chromosomes from *sycp1*^+/+^; *dmrt1*^−/−^ (A) and *sycp1*^−/−^; *dmrt1*^−/−^ (B) oocytes stained for Sycp3 (green) and Sycp1 (magenta and gray). Examples of spread chromosomes from meiotic prophase described in Fig 3 A–B. Scale bar = 10 µm.

**Supplementary Figure S4.** yH2AX staining in *sycp1^−/−^* testes and oocytes

(A) Whole mount testes of *sycp1*^+/+^ and *sycp1*^−/−^ males stained with DAPI and γH2AX (green). Dashed lines represent metaphase nuclei. Examples of spermatocytes in metaphase (MI) and sperm are noted. Scale bar = 50 µm.

(B) Whole mount ovaries of *sycp1*^+/+^ and *sycp1*^−/−^ males stained with DAPI and γH2AX (green). Examples of Stage IA oocytes (leptotene/zygotene stage) and Stage IB oocytes are indicated by blue lines. Scale bar = 50 µm.

**Supplementary Figure S5.** Model of chromosome events during meiotic prophase.

(A–C) Temporal progression of chromosome axis formation, DSB repair, Hormad1 localization, synapsis and homolog juxtaposition. (A) In wild-type zebrafish, DSBs form in early prophase and are gradually repaired. Hormad1 localizes to asynapsed regions and is displaced upon synapsis. As synapsis progresses, homologs are “zippered up”. (B) In *syce2* mutants, there is partial recruitment of Sycp1 to synapsed-like regions and unaligned axes. Hormad1 localizes along the entire length of chromosomes. DSBs form but their repair is less efficient. Without a fully formed SC, chromosome co-alignment is limited to sub-telomeric regions. (C) In *sycp1* mutants, there is no synapsis, Hormad1 localizes along entire axes and DSB repair is less efficient. Chromosome co-alignment is also limited to sub-telomeric regions.

**Supplementary File 1.** Master data table with graph values

## Notes

### Competing Interest Statement

The authors have declared no competing interest.

