## Supplementary figures and images for "Zebrafish lack a strong meiotic checkpoint response to defects in chromosome synapsis"

### Supplemental Figure 1

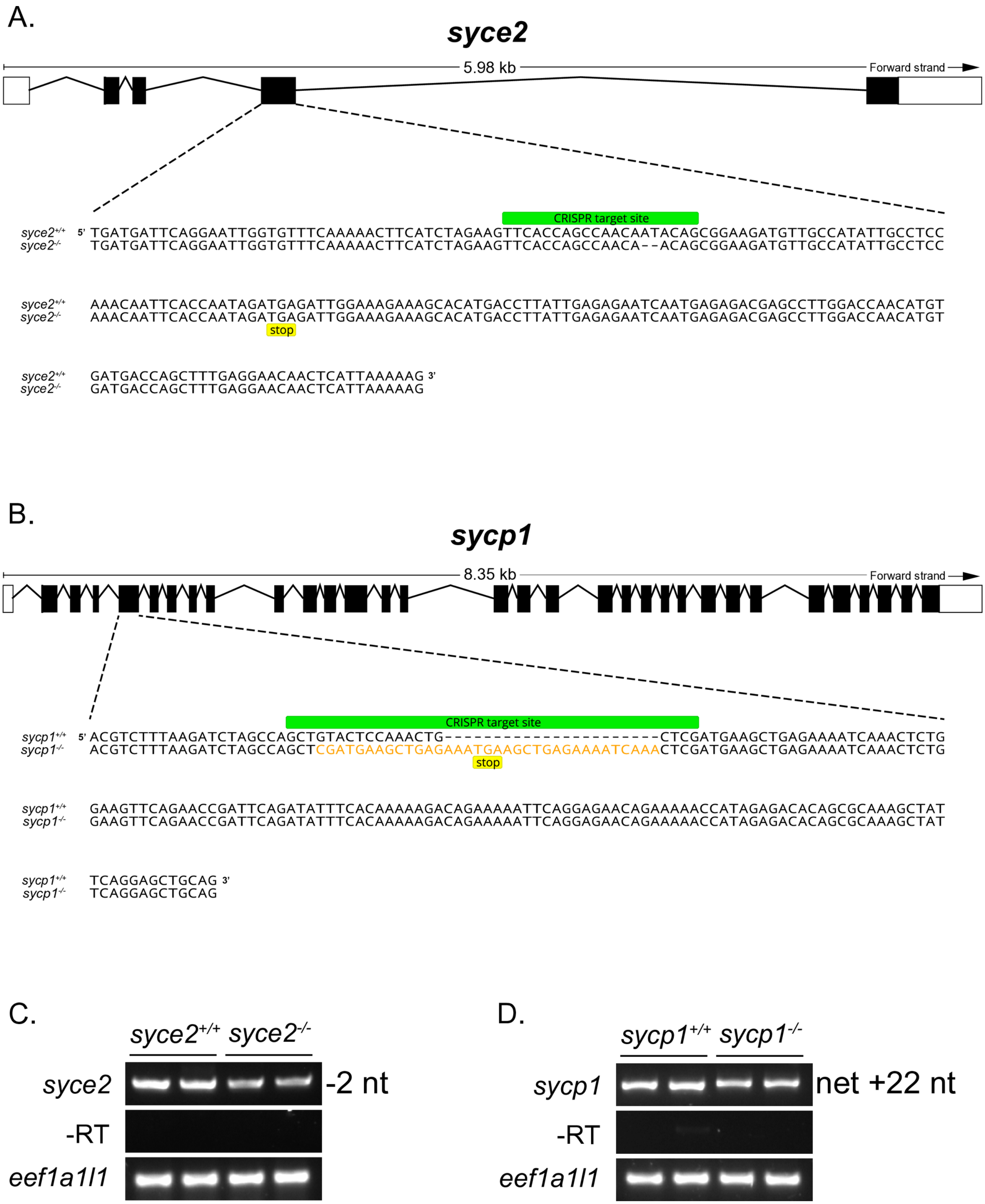

### Supplemental Figure 2

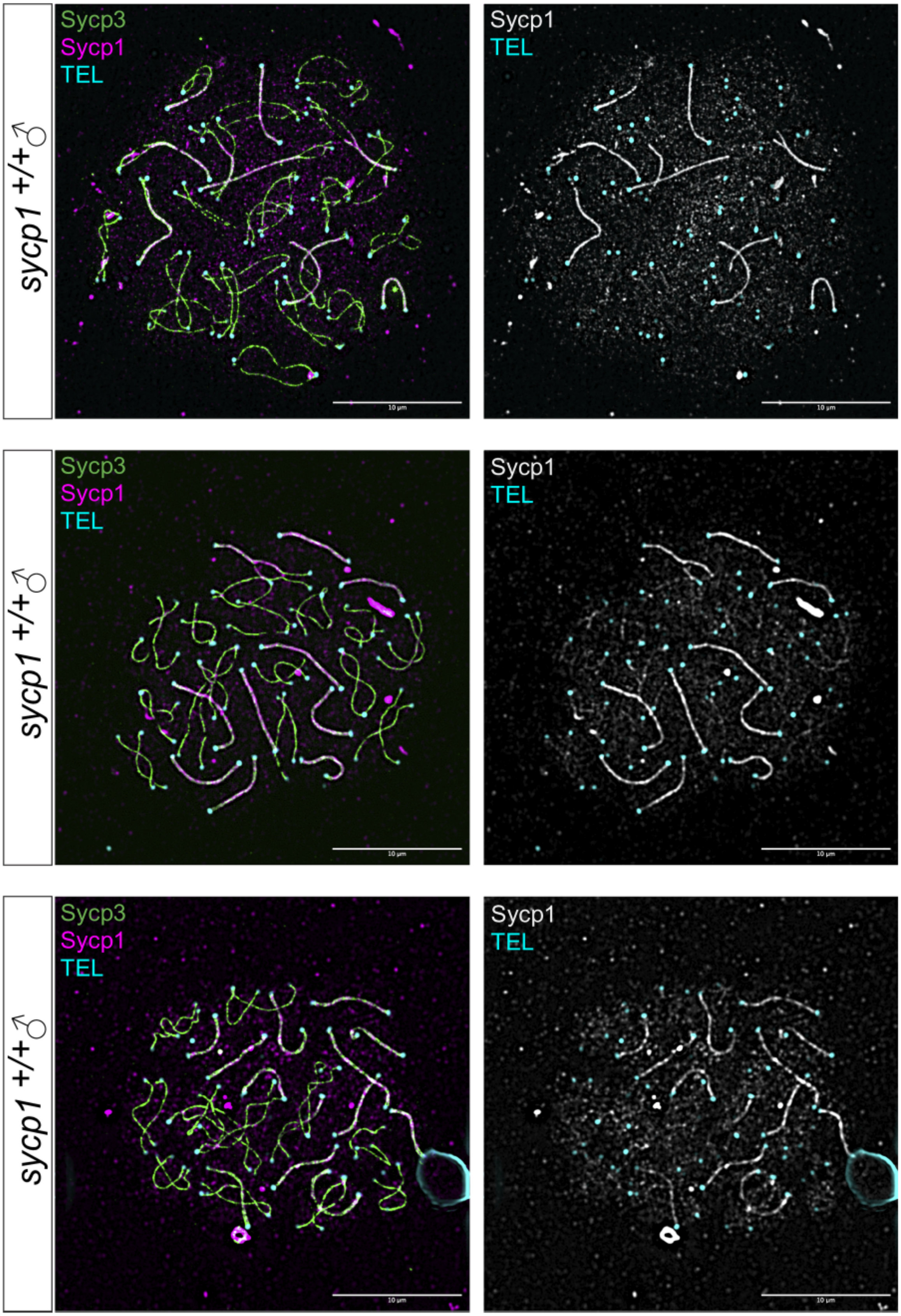

### Supplemental Figure 3

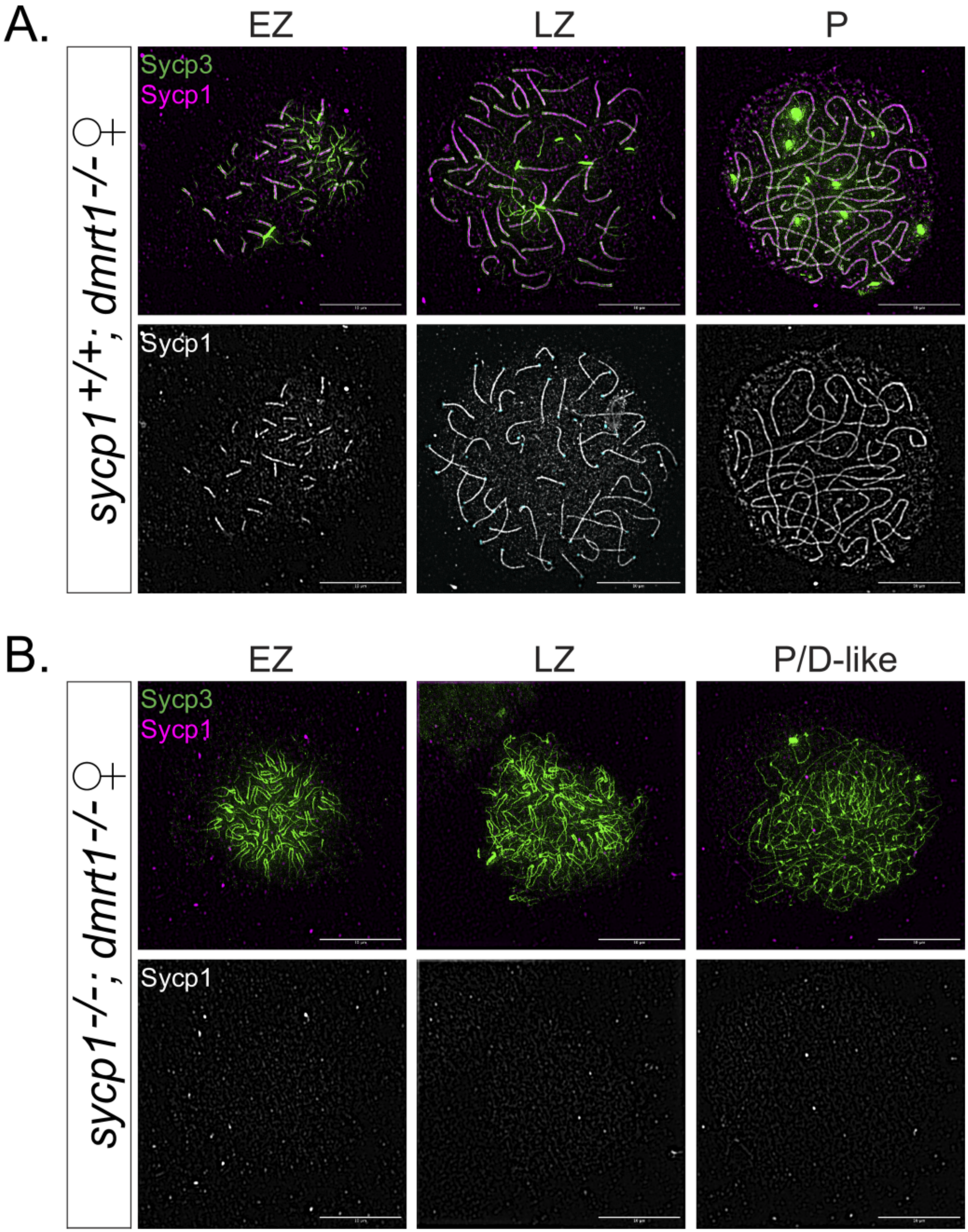

### Supplemental Figure 4

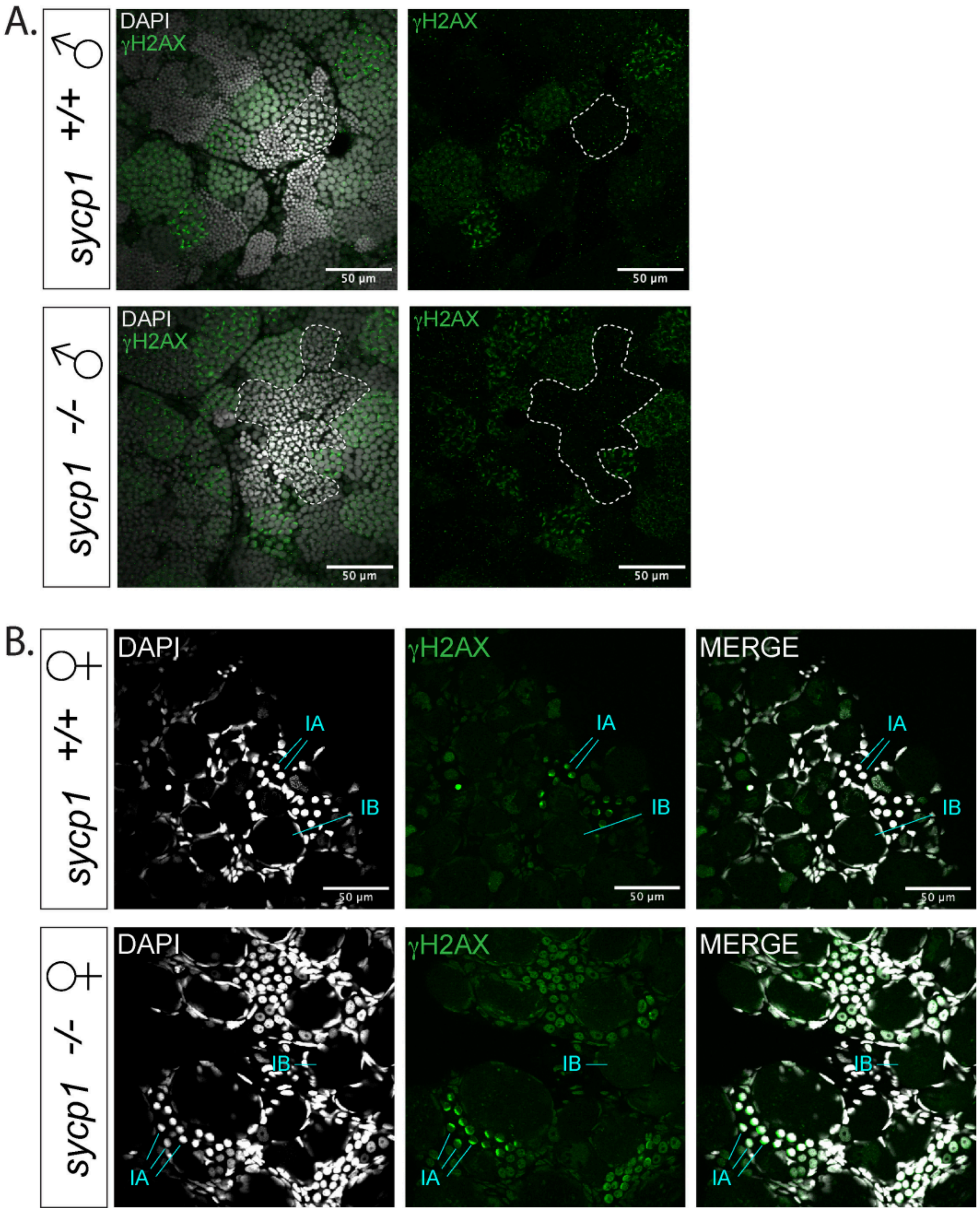

### Supplemental Figure 5

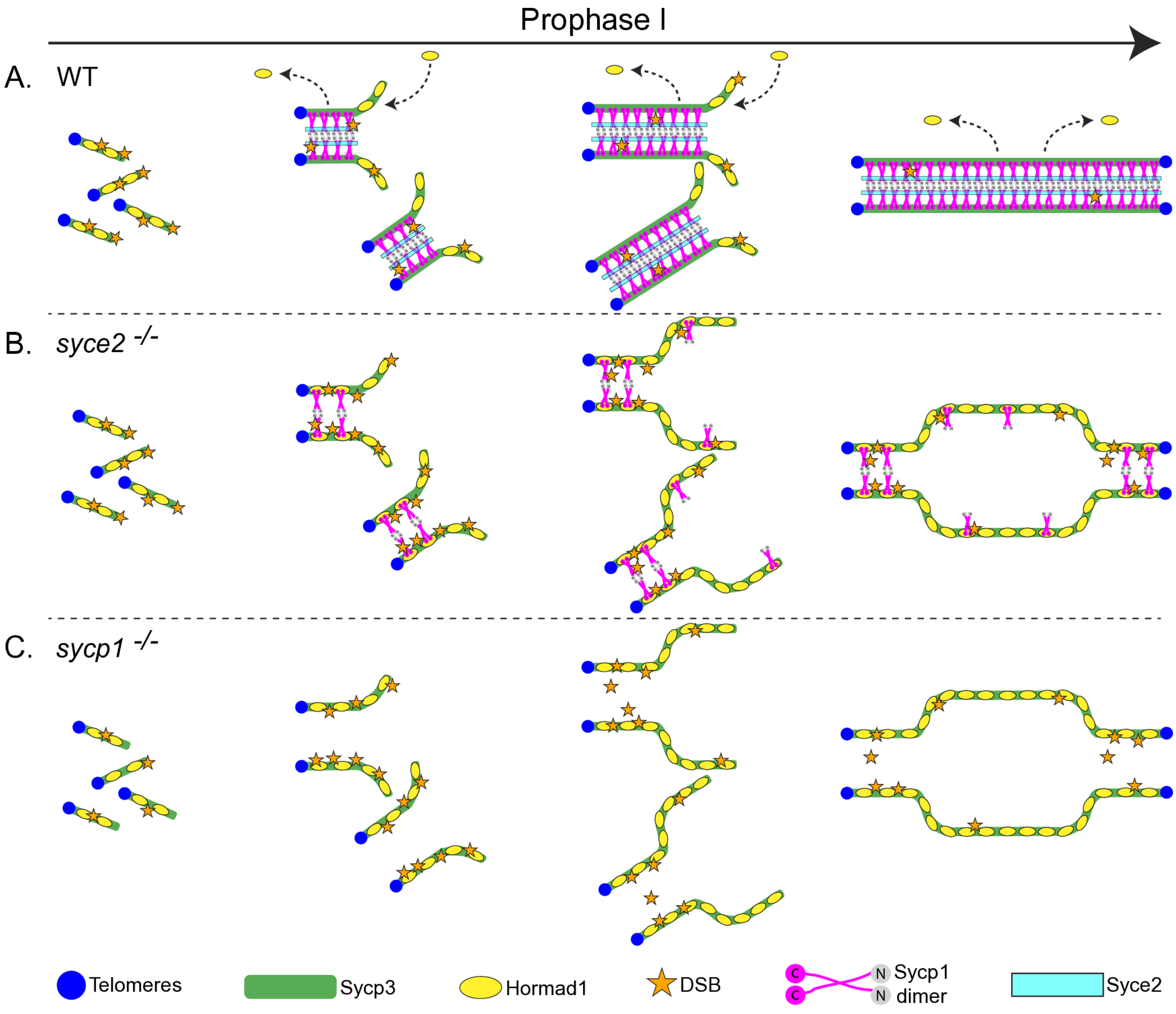
